# Harmoni: a Method for Eliminating Spurious Interactions due to the Harmonic Components in Neuronal Data

**DOI:** 10.1101/2021.10.06.463319

**Authors:** Mina Jamshidi Idaji, Juanli Zhang, Tilman Stephani, Guido Nolte, Klaus-Robert Müller, Arno Villringer, Vadim V. Nikulin

## Abstract

Cross-frequency synchronization (CFS) has been proposed as a mechanism for integrating spatially and spectrally distributed information in the brain. However, investigating CFS in Magneto- and Electroencephalography (MEG/EEG) is hampered by the presence of spurious neuronal interactions due to the non-sinusoidal waveshape of brain oscillations. Such waveshape gives rise to the presence of oscillatory harmonics mimicking genuine neuronal oscillations. Until recently, however, there has been no methodology for removing these harmonics from neuronal data. In order to address this long-standing challenge, we introduce a novel method (called HARMOnic miNImization - Harmoni) that removes the signal components which can be harmonics of a non-sinusoidal signal. Harmoni’s working principle is based on the presence of CFS between harmonic components and the fundamental component of a non-sinusoidal signal. We extensively tested Harmoni in realistic EEG simulations. The simulated couplings between the source signals represented genuine and spurious CFS and within-frequency phase synchronization. Using diverse evaluation criteria, including ROC analyses, we showed that the within- and cross-frequency spurious interactions are suppressed significantly, while the genuine activities are not affected. Additionally, we applied Harmoni to real resting-state EEG data revealing intricate remote connectivity patterns which are usually masked by the spurious connections. Given the ubiquity of non-sinusoidal neuronal oscillations in electrophysiological recordings, Harmoni is expected to facilitate novel insights into genuine neuronal interactions in various research fields, and can also serve as a steppingstone towards the development of further signal processing methods aiming at refining within- and cross-frequency synchronization in electrophysiological recordings.

## 1. Introduction

The importance of oscillatory neuronal activity has been demonstrated by its association with cognitive, sensory, and motor processes in the brain (Buzsáki and Draguhn, 2004; Engel and Fries, 2010; Harris and Gordon, 2015; Miller et al., 2010; Sadaghiani and Kleinschmidt, 2016). Various oscillatory processes have to be integrated in order to support formation of behaviorally relevant outputs based on a multitude of sensory and cognitive factors. This neuronal integration is facilitated by complex spatial connectivity patterns in the brain (Bullmore and Sporns, 2009; Nentwich et al., 2020). In this context, phase-phase synchronization (PPS) has been hypothesized to represent a mechanism through which such spatially distributed information can be integrated in the brain with a high temporal precision (Fries, 2015). Importantly, PPS underlies not only spatially, but also spectrally distributed interactions - so-called cross-frequency synchronization (CFS) (Canolty and Knight, 2010; Jensen and Colgin, 2007; Nikulin and Brismar, 2006; Palva et al., 2005; Palva and Palva, 2018a,b). Magneto- and Electroencephalography (MEG/EEG) provide a unique opportunity to non-invasively study these neuronal interactions in humans.

Since in the frequency domain analysis the kernel function is sinusoidal, we often conceptualize oscillations as sinusoids. However, neural oscillations with non-sinusoidal waveshape are abundant in human electrophysiological recordings Cole and Voytek (2017). Such non-sinusoidality reflects complex trans-membrane ion currents flowing though highly morphologically asymmetric neurons (e.g. pyramidal cells) where inward and outward currents are unlikely to balance each other with the exact temporal dynamics thus leading to different shape of oscillations recorded with EEG/MEG/LFP (Local field potential) (Jones et al., 2009). This ubiquity of the non-sinusoidal waveform of brain oscillations has significant implications for the analysis of brain connectivity.

A periodic signal can be decomposed into its harmonic components using Fourier analysis. For the sake of clarity, we call the first harmonic the fundamental component and from here on by *harmonics* we mean the second and higher harmonic components whose central frequencies are integer multiples of the fundamental frequency. By band-pass filtering the signal around the fundamental and harmonic frequencies, we can separate the respective components, which are – by construction – CF synchronized to the fundamental component (Hyafil, 2017; Scheffer-Teixeira and Tort, 2016). Additionally, if the band-pass filters of the harmonics frequency are wide enough, a phaseamplitude coupling (PAC) can be observed between the fundamental and harmonic components (Giehl et al., 2021; Hyafil, 2017). Note that, as also discussed in (Kramer et al., 2008), non-sinusoidal signals can be constructed from the mixture of distinct sources with cross-frequency coupling. However, in this work, we do not distinguish whether the non-sinusoidality originates from signal mixing or the intrinsic waveshape of the signal. In the discussion section, we elaborate on the effect of signal mixing.

In this manuscript, we address the effects of non-sinusoidal shape of the brain oscillations on the observation of spurious interactions between the oscillatory brain activities. In spite of other spurious interactions (e.g. bias of the data length), the spurious interactions due to the waveshape cannot be determined by statistical methods. For example, our recently introduced method for separating cross-frequency coupled sources cannot distinguish sources with genuine interactions and those which are coupled because of the higher frequency signal being the harmonic of the lower frequency one (Idaji et al., 2020) because a harmonic-driven synchronization is not statistically distinguishable from a genuine coupling. Therefore, distinguishing harmonic-driven and genuine interactions has currently gained more attention and still remains as a major challenge in the MEG/EEG connectivity research (Giehl et al., 2021; Scheffer-Teixeira and Tort, 2016; Siebenhühner et al., 2020). The main reason of this challenge is that the connectivity analysis of MEG/EEG data is typically done using band-pass filtering, which separates the fundamental and harmonic components of an oscillatory activity with a non-sinusoidal waveform. As a result, the observed within- and cross-frequency synchronization between the components in the frequency bands of the fundamental and harmonic frequencies can be mistakenly interpreted as genuine interaction. Figure 1 shows a schematic example where two non-sinusoidal signals are synchronized. This coupling should be manifested in the synchronization of the fundamental components, while the harmonic components shape the waveform of the individual signals. However, the harmonic components are also spuriously synchronized and additional CFS is observed between and within the regions. Since these interactions (shown in dashed lines in figure 1-B) are observed due to the waveform of the individual signals, they are referred to as spurious, in contrast to genuine interactions. The omnipresence of these spurious interactions in all human MEG/EEG recordings makes the validity of the previously studied within- and crossfrequency connectivity maps ambiguous.

**Figure 1:**
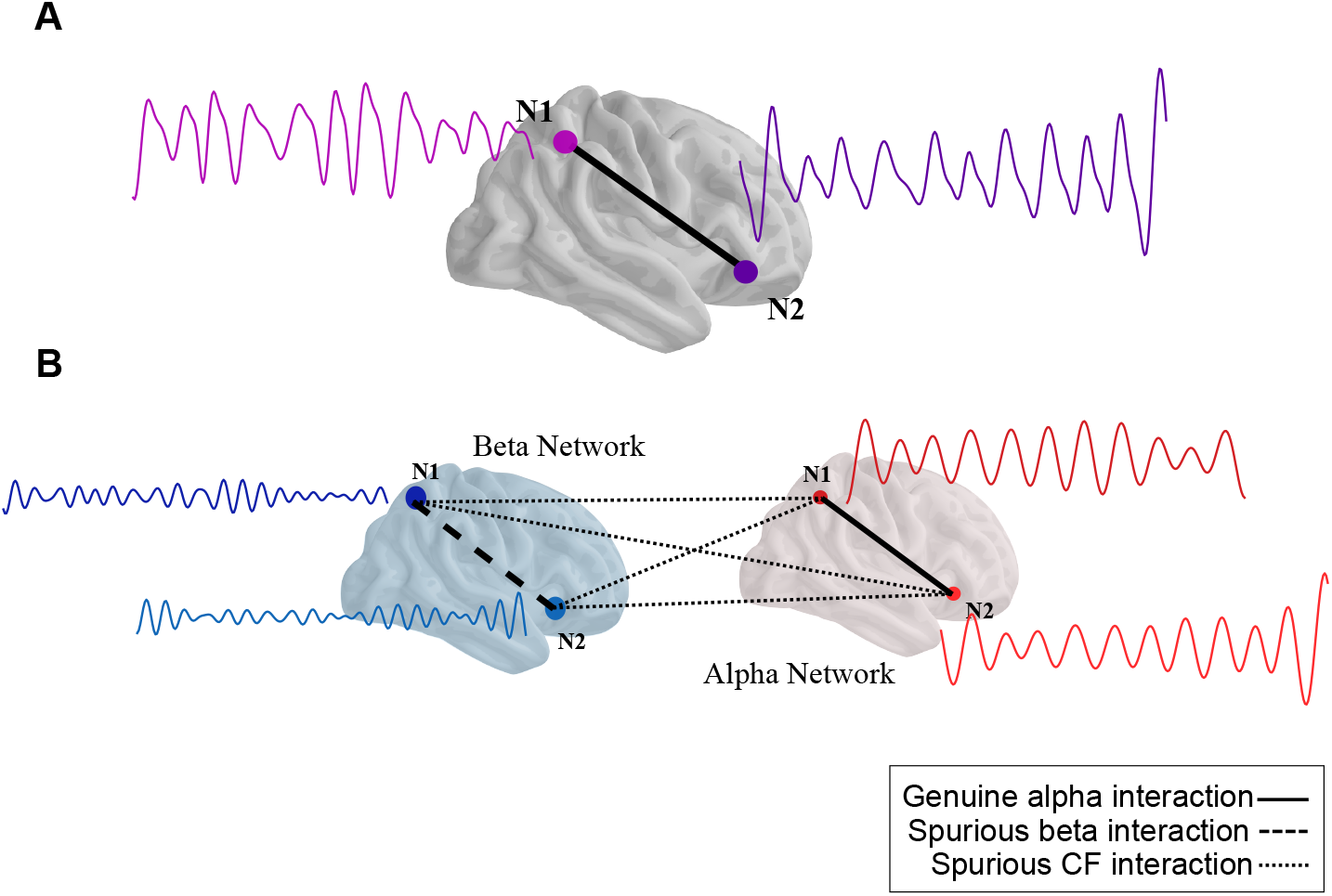
How non-sinusoidal shape of the neuronal oscillations impacts the connectivity of brain regions. Panel A shows two non-sinusoidal oscillations with their fundamental frequency in the alpha band. The second harmonics of these signals are located in the beta band. As a byproduct of the coupling of the fundamental alpha components (the solid line in panel B), the second harmonics are also coupled to each other, which results in spurious interactions within the beta band (the dashed line in panel B) and across the two frequency bands (dotted lines in panel B).

There has been an attempt from Siebenhühner et al. (2020) to *discard* the potentially spurious connections from cross-frequency (CF) connectivity graphs based on the detection of ambiguous motifs in the connectivity graphs. In that work, any CFS connection forming a triangle motif with the local CFS and within-frequency inter-areal phase synchronization is considered as ambiguous and is discarded. However, such an approach cannot disentangle the within-frequency spurious interactions in the harmonic frequency bands, and is specific to the CF connectivity graphs. Furthermore, this approach cannot distinguish cases of genuine couplings which form an ambiguous motif. A more attractive approach, however, would remove or suppress the data components that can be associated with the harmonics of the periodic neuronal activity. Such an approach can provide the opportunity of using the cleaned narrow-band data (in the frequency range of the harmonics) for within-frequency and cross-frequency connectivity analyses.

In the current work, we introduce a novel, first-of-its-kind method for removing effects of harmonics on the estimation of within- and cross-frequency synchronization. Our method, called HARMonic miNImization (Harmoni), is (to the best of our knowledge) the first existing signal processing tool for suppressing higher harmonic components of a periodic signal, without band-stop filtering or rejecting non-sinusoidally shaped signal components using ICA or any other multi-variate decomposition.

We extensively tested Harmoni with realistic EEG simulations and show that the spurious interactions are alleviated significantly, while the genuine activities are not affected. Harmoni is then applied to resting-state EEG (rsEEG) data and we show that the CFS connections mimicking genuine interactions are suppressed, while many masked remote interactions are recovered.

## 2. Materials and Methods

### 2.1. Phase-Phase Synchronization

Phase-Phase Synchronization (PPS) can be defined for within-frequency as well as for cross-frequency (CF) interactions. In order to define the within- and cross-frequency synchronization indices, assume two complex narrow-band signals 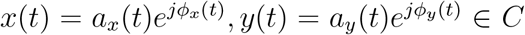 with central frequencies *f_x_* and *f_y_*, respectively. Here, by *narrow-band complex signal* we mean the analytic signal built using the Hilbert transform. Formally, if *x_H_* (*t*) is the Hilbert transform of a narrow-band real signal *x_R_*(*t*) = *a_x_*(*t*) cos (*ϕ_x_*(*t*)), then *x*(*t*) = *x_R_*(*t*) + *jx_H_* (*t*) is the analytic signal of *x_R_*(*t*). In these formulations the index *R* indicates that the signal is real valued and the index *H* denotes a Hilbert transformed signal. Note that, another way to get the narrow-band complex signals from a broad-band signal is complex wavelet transforms.

If *f_x_* = *f_y_* then *x*(*t*) and *y*(*t*) are two narrow-band signals in the same frequency band. Their complex-valued coherence *coh*(*x, y*) ∈ *C* can be computed from the following equation:

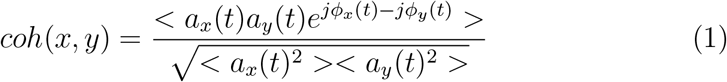

where *< . >* is the averaging operator over time and 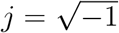 is the imaginary number.

We use the absolute of the imaginary part of coherence (iCoh) (Nolte et al., 2004) for estimating the connectivity between two signals in the same frequency band. This prevents a lot of the within-frequency spurious interactions due to signal mixing and volume conduction in EEG.

If *nf_x_* = *mf_y_* for *m, n ∈ N*, the cross-frequency synchronization (CFS, known as m:n synchronization) of *x*(*t*) and *y*(*t*) can be quantified by m:n absolute coherence *coh_m_*_:*n*_(*x, y*) ∈ *R* defined by the following equation:

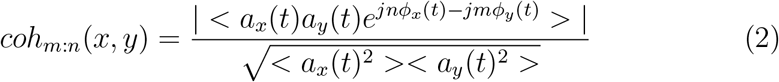

which is in principle similar to m:n phase locking value as:

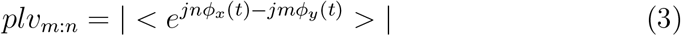

with the difference that in equation 2 the amplitudes of the signals are taken into account and the phase estimations during higher amplitudes are weighted higher. Giehl et al. (2021) have used a variant of equation 2. Equation 2 reduces to the absolute part of equation 1 for *m* = *n* = 1. In this work, we are specifically interested in the case that *m* = 1 and *n* > 1, i.e. when *x*(*t*) is a signal with central frequency *f_x_* and *y*(*t*) is a faster oscillation with the central frequency *f_y_* = *nf_x_*. In this case, *coh*_1:*n*_(*x, y*) = |*coh*(*x_n_, y*)|, where 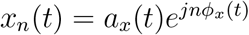 is built by multiplying the phase of *x*(*t*) by *n*, i.e. accelerating *x*(*t*) by a factor of *n*.

CFS as defined by equation 2 has a real value between 0 and 1, with 0 corresponding to the lack of any phase synchronization between two completely independent signals and 1 for two perfectly synchronized time-series with the same amplitude envelope.

### 2.2. Genuine vs. spurious interactions

The PPS and CFS indices of equations 1 and 2 have a bias based on the length of the data time-series, i.e., two band-pass filtered random time-series also show a value larger than 0. Therefore, a test of significance is necessary for phase synchronization measures (Scheffer-Teixeira and Tort, 2016) in order to distinguish such spurious interactions when the data length is not sufficient.

Another type of spurious interactions (which is not statistically discernible from real interactions) is the interactions due to the waveshape of brain signals. The reason is that harmonic components of a signal with a non-sinusoidal shape have CFS to each other. As an illustrative example, figure 2 depicts a sawtooth-shaped signal and its fundamental and 7th harmonic components. The 7th harmonic of this sawtooth-shaped signal has an almost perfect 1:7 synchronization to the fundamental frequency (*coh*_1:7_ = 0.99). Additionally, although it is not the focus of this manuscript, it is interesting to note that when a non-sinusoidally shaped signal (here sawtooth-shaped) is filtered in a wider frequency range around the harmonic frequency, PAC is observed between the harmonic and fundamental frequencies (in addition to CFS). In this paper, however, our focus is on the n:m synchronizations.

**Figure 2:**
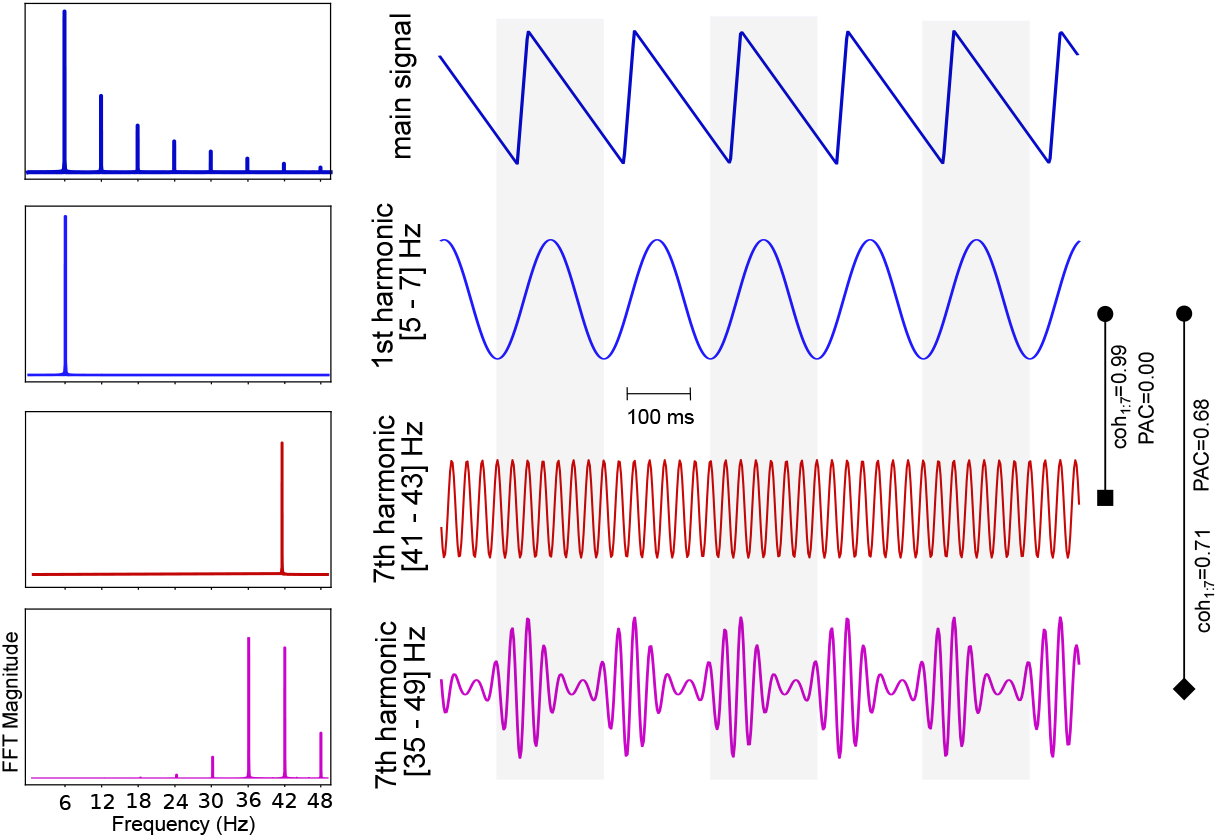
A simulated sawtooth-shaped signal with the fundamental frequency equal to 6 Hz is depicted in the first row and the fundamental 6 Hz component (i.e. the 1st harmonic) is shown in the second row. The 7th harmonic component filtered at a frequency window with width of 2Hz is illustrated in third row. Additionally, the sawtooth signal was filtered around the 7th harmonic frequency with a window size of 7Hz, depicted in the fourth row. The magnitude of the fast Fourier transform (FFT) of each signal is depicted at its left side. The CFS and PAC between the fundamental component and the two components with central frequency of the 7th harmonic frequency are noted along the right side vertical lines. The 7th harmonic on the third row shows a strong 1:7 synchronization to the fundamental component (*coh*_1:7_ = 0.99) and no PAC. However, if filtered at a wider frequency band, the harmonic component shown on the fourth row shows also a PAC with the fundamental component. Note that the amplitude of the signals and their FFT magnitudes are scaled arbitrarily for the sake of better illustration.

The example of figure 2 shows that by band-pass filtering a single process one can observe cross-frequency coupling between its different components, although these components still represent the same complex signal. In the literature of cross-frequency coupling (Hyafil, 2017; Scheffer-Teixeira and Tort, 2016; Siebenhühner et al., 2020; Giehl et al., 2021), such a coupling between the components of a single process, or generally an interaction between two signals where *at least one of them* is a higher harmonic of a non-sinusoidal process is called *spurious*. This is usually in contrast to *genuine* interactions between two signals representing two distinct processes where *none of them* is a higher harmonic of a periodic signal. Formally, let 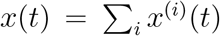 and 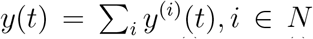 be two n:m synchronized periodic oscillatory processes, where *x*^(*i*)^ and *y*^(*i*)^ are the i-th harmonic components of *x*(*t*) and *y*(*t*), respectively. The fundamental components (*x*^(1)^ and *y*^(1)^) and higher harmonics (*x*^(*i*)^ and *y*^(*i*)^ for *i* > 2) of each of these signals can be separated from each other by band-pass filtering *x*(*t*) and *y*(*t*). The synchronization of *x* and *y* implies that for any 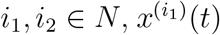 and 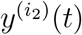 are cross-frequency synchronized. When assessing the synchronization of the narrow-band signals, we consider only the synchronization of fundamental components *x*^(1)^ and *y*^(1)^ genuine. The synchronization of 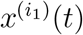 and 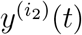 for *i*_1_ > 1 or *i*_2_ > 1 is *harmonic-driven* and is called spurious. Note that this does not mean that the signal components are not synchronzed and the synchronization value is non-zero because of insufficient number of data points or due to filtering. By spurious interactions due to waveshape it is meant that any coupling including higher harmonics is in fact mediated by the fundamental component of the respective non-sinusoidal signal. Figure 1 illustrates various possible within- and cross-frequency spurious synchronizations due to waveshape. In the next section we introduce an original signal processing method for suppressing the harmonic-driven synchronizations in connectivity analyses using electrophysiological data.

A final important note is that, as discussed in (Kramer et al., 2008), a non-sinusoidal signal can be constructed from the mixing of distinct sources with CFS or PAC. This is actually a major concern in electrophysiological research even outside of connectivity topic. Although we do not account for this issue in our analyses explicitly, we discuss it in the discussion section, “Harmoni and signal mixing”.

### 2.3. HARMOnic miNImization (HARMONI)

Assume that *z*(*t*) = *s*(*t*) + *ϵ*(*t*), where *s*(*t*) is a a periodic signal with the fundamental frequency of *f*_0_. *ϵ*(*t*) is additive noise or any other process such as another oscillatory activity mixed with *s*(*t*). Harmoni aims at removing the components of *z*(*t*) within a narrow frequency band around *nf*_0_, *n ∈ N, n* ≥ 2 that have similar phase profile as the fundamental component of *s*(*t*). For this purpose, we can write *z*(*t*) = *x_R_*(*t*) + *y_R_*(*t*) + *ξ*(*t*), where *x_R_*(*t*) = *a_x_*(*t*) cos (*ϕ_x_*(*t*)) and *y_R_*(*t*) = *a_y_*(*t*) cos (*ϕ_y_*(*t*)) are the real-valued contents (indicated by the index *R*) from frequency bands *f*_0_ and *nf*_0_, respectively. *ξ*(*t*) represents all other components of *z*(*t*) except *x_R_*(*t*) and *y_R_*(*t*). Therefore, *x_R_*(*t*) and *y_R_*(*t*) are estimated using band-pass filtering *z*(*t*) within the respective frequency bands of the fundamental and harmonic frequencies. We define *x*(*t*) and *y*(*t*) as the analytical signals of *x_R_*(*t*) and *y_R_*(*t*) built using the Hilbert transform and work with them in the next steps of Harmoni. Note that *x*(*t*) and *y*(*t*) can be also generated by applying complex wavelet transforms to *z*(*t*).

The fundamental component of a non-sinusoidal signal has 1:n synchronization to its n-th harmonic component. Therefore, the phase information of the harmonic components can be recovered from the phase of the fundamental component. Using *x*(*t*), Harmoni tries to remove the parts of *y*(*t*) that are 1:n coupled to *x*(*t*), or equivalently 1:1 coupled to 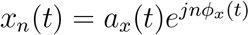.

As mentioned above, the part of *y*(*t*) which is a harmonic of a component in *x*(*t*) should be phase synchronized to *x_n_*(*t*). Therefore, we estimate the harmonic part of *y* by *λx_n_*(*t*), *λ* ∈ *C*. *y_corr_*(*t*) = *y*(*t*) − *λx_n_*(*t*) contains the non-harmonic components of *y*(*t*), where *y_corr_*(*t*) has a minimum possible within-frequency synchronization to *x_n_*(*t*). The complex multiplier *λ* = *ce^jϕ^* is estimated through the following optimization problem:

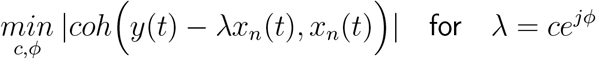

Here, the phase of *λ* compensates the possible phase difference between the harmonic and fundamental components. Figure 3 shows a schematic block diagram of Harmoni. Practically, we perform a grid-search procedure for computing *λ* = *ce^jϕ^*, which is presented in algorithm 1. In practice, in a connectivity pipeline, the activity of each brain site - that can be a region-of-interest (ROI) or an electrode - is band-pass filtered within the two bands of interest, namely *f*_0_ and *nf*_0_. Then Harmoni is applied on the data of each sensor or ROI. In the next section, it will be described in detail how Harmoni can be used in a connectivity analysis pipeline with electrophysiological data.

**Figure 3:**
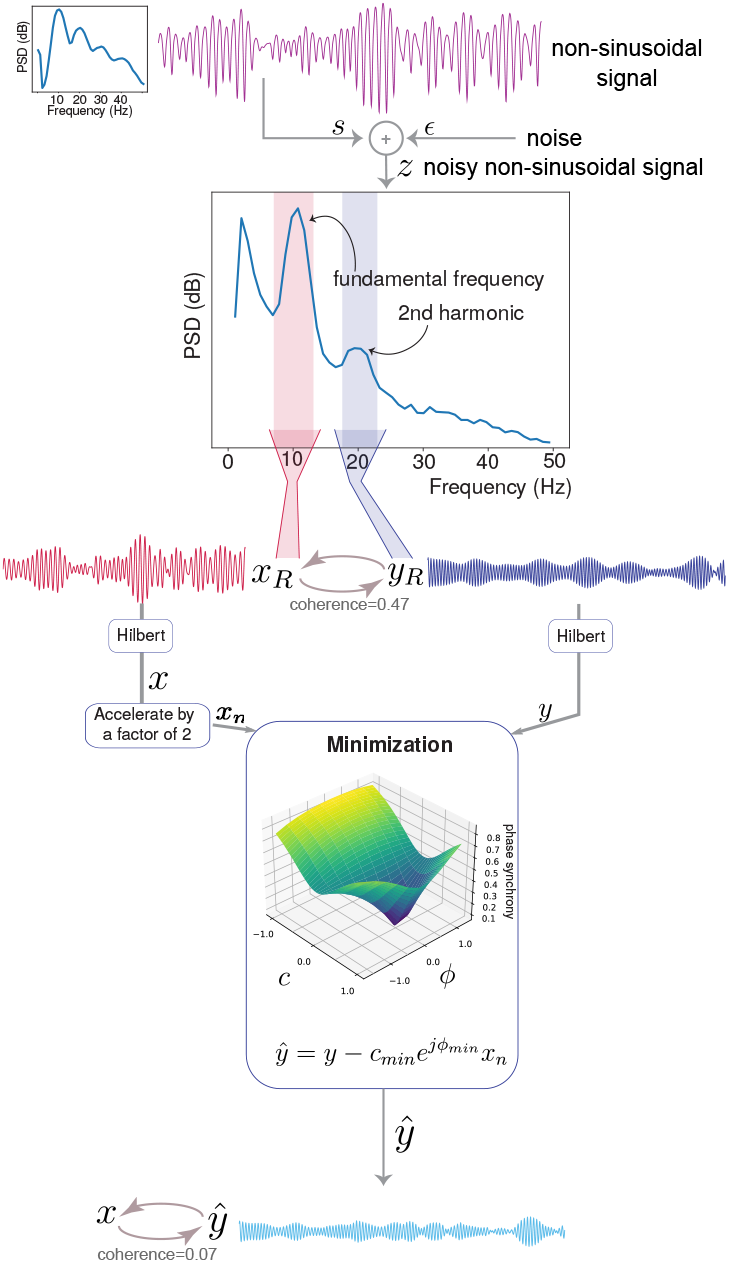
Harmoni is a method that removes harmonics of a non-sinusoidal signal. The inputs are the band-pass filtered signals in the frequency bands of the fundamental and harmonic frequencies. In this figure, the signal is a non-sinusoidal alpha rhythm with fundamental and second harmonic frequencies of 10Hz and 20Hz, respectively. The band-pass filtered signals at 10Hz and 20Hz are used as inputs to the minimization block, which runs a regression-like algorithm to find the best multiplier for removing the harmonic parts of *y*(*t*). This is done by means of subtracting a scaled version of *x_n_*(*t*) from *y*(*t*), where *x_n_*(*t*) is an accelerated version of *x*(*t*) by multiplying its phase by a factor of *n* (here *n* = 2). The output of Harmoni is a band-limited signal in the harmonic frequency band (here 20Hz - the second harmonic) where the harmonic component is removed.

### 2.4. Connectivity pipeline in source space

Figure 4 shows a block-diagram of a connectivity pipeline, also implementing Harmoni. The first step is to band-pass filter the multi-channel data within the frequency bands of interest *f*_0_ and *nf*_0_. For instance, if we are interested in alpha and beta band, *f*_0_ = 10 and *n* = 2. Below, we will elaborate upon the next steps.

**Figure 4:**
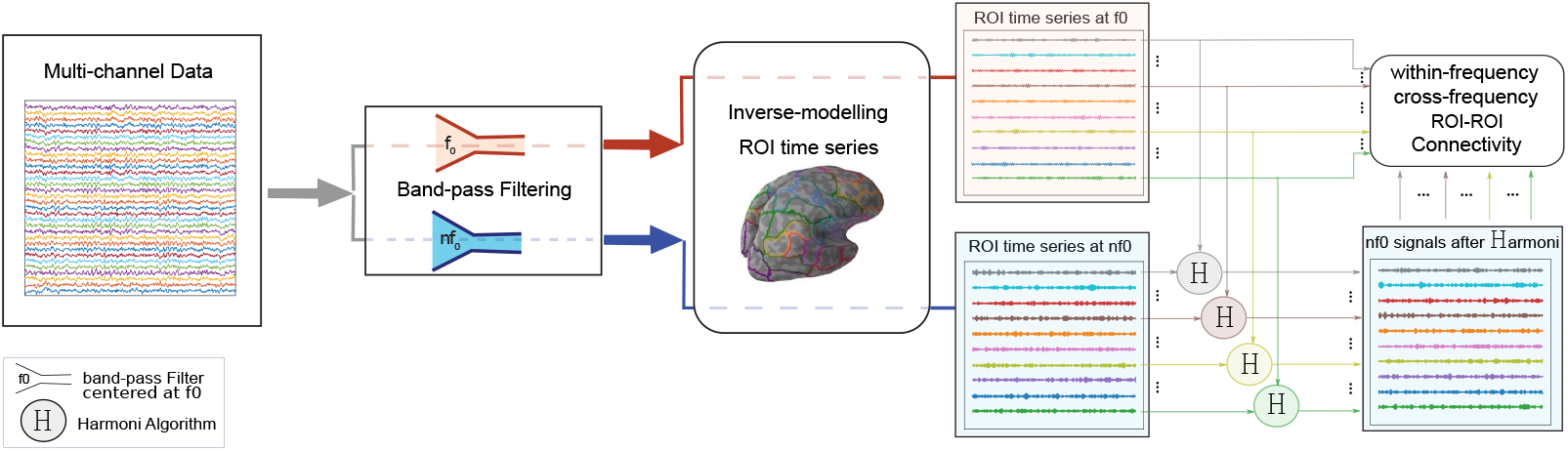
The block-diagram of Harmoni pipeline in source space. The multi-channel signal is first band-pass filtered in the range of the fundamental frequency (*f*_0_) and the harmonic frequency of interest (*nf*_0_). The narrow-band signals are mapped to the cortical surface using the inverse solution and the ROI time series are extracted. The ROI signals in the range of harmonic-frequency are then corrected with Harmoni and the potential harmonic components are removed. Finally, the ROI-ROI within- and cross-frequency connectivity maps are computed. In this paper, without loss of generality and due to the better SNRs, we set *f*_0_ = 10 and *n* = 2.

#### 2.4.1. Forward and inverse solutions

We used fsaverage standard head model and the three-layer boundary element model (BEM) accompanied with MNE Python (Gramfort et al., 2013, 2014). 64 electrodes (or a subset of it) with positions according to the BioSemi cap were used and aligned to the MRI coordinates. MNE-Python was used to create a dipole grid on white matter surface with oct6 spacing between the grid points, resulting in 4098 sources per hemisphere. The surface-based source space and the BEM solutions were then used for computing a forward solution. An inverse solution with dipole directions normal to the cortical surface was computed with eLORETA inverse modelling (Pascual-Marqui, 2007) with the regularization parameter equal to 0.05, and the noise covariance equal to the covariance of 64 white-Gaussian signals with equal duration to the data, which is an estimation of the identity matrix.

#### 2.4.2. From sensor space to ROIs

The band-pass filtered multi-channel EEG data were projected to the cortical surface using the computed inverse solution, resulting in ∼8000 reconstructed surface sources. These sources were then grouped based on an atlas into regions of interest (ROIs). We used the Desikan Killiany atlas with 68 ROIs (Desikan et al., 2006) for simulations and Schaefer atlas with 100 ROIs (Schaefer et al., 2018) for real data analysis. Singular value decomposition (SVD) was then applied to the band-pass filtered time-series of the sources of each ROI and a single time-series was computed per ROI. As a result, the ∼8000 reconstructed cortical sources were translated to *n_ROI_* ROI times-series in each frequency band (here: *n_ROI_*=number of ROIs in the used atlas), which are ready for connectivity computations.

#### 2.4.3. Harmoni

Although the ROI time series can be directly used for computing the connectivity maps, we suggest to use Harmoni as an intermediate step in a connectivity pipeline. Harmoni is applied on the signals of each ROI in the two frequency bands of interest centered at *f*_0_ and *nf*_0_, which correspond to the fundamental and the n-th harmonic frequencies. The output of the algorithm is a signal in the frequency band of *nf*_0_ for which the harmonic components are suppressed to a large extent. The ROI time series at *f*_0_ and the Harmoni-corrected signals at *nf*_0_ are then passed to the next step for computing the within- and cross-frequency synchronization maps.

#### 2.4.4. From ROIs’ time-series to connectivity maps

For both of the simulations and real data, after computing the ROI time series and applying Harmoni on them, we computed a connectivity index for each pair of the ROIs, resulting in an *n_ROI_ × n_ROI_* graph. For within-frequency connectivity (here in alpha and beta bands), we used the absolute of imaginary part of coherence (iCoh) computed from the imaginary part of equation 1 and for the cross-frequency synchronization we used the extension of coherence for n:m coupling as in equation 2.

### 2.5. Simulations

#### 2.5.1. Signals and SNR

The pipeline for producing signals and the definition of signal-to-noise ratio (SNR) are similar to that of (Idaji et al., 2020). In this section we describe the procedure of simulating the signals and how SNR is defined in our simulation pipelines. Note that in all places, band-pass filtering was carried out using fourth-ordered Butterworth filters designed for the frequency band of interest. The filtering was applied forward and backward in order to avoid phase shift in data.

##### Additive noise

The time-series of the noise sources were produced with the colornoise package (Patzelt, 2019) in Python by building a random signal with a 1/f (pink) spectrum from a random white Gaussian noise.

##### Sinusoidal oscillations

Without loss of generality, in our simulations, all of the time-series of the sinusoidal oscillatory sources were simulated in alpha (8-12 Hz) and beta (16-24 Hz) frequency bands.

Independent sources (those without a synchronization to other source signals) were generated by band-pass filtering white Gaussian noise in the frequency band of interest. The analytic signals of these oscillations were built using the Hilbert transform of them. For instance, if *x_R_*(*t*) is an alpha oscillation produced by band-pass filtering white Gaussian noise within (8-12) Hz and *x_H_* (*t*) is the Hilbert transform of *x_R_*(*t*), *x*(*t*) = *x_R_*(*t*) + *jx_H_* (*t*) is the analytic signal of *x_R_*(*t*).

A source signal *y*(*t*) with 1:n synchronization to an oscillation *x*(*t*) was simulated by phase-warping of *x*(*t*), i.e.:

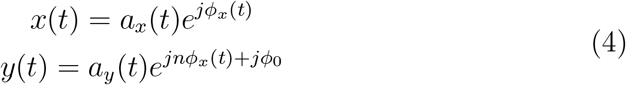

where *x*(*t*) ∈ *C* is the analytic signal of an oscillation generated by band-pass filtering white Gaussian noise around *f*_0_, *y*(*t*) ∈ *C* is the analytic signal of an oscillation within a frequency band around *nf*_0_ and 1:n synchronized to *x*(*t*), and *ϕ*_0_ is the phase difference of the two signals taken randomly from a uniform distribution between [− *π/*2, *π/*2]. *a_y_*(*t*) is either equal to *a_x_*(*t*) or equal to the envelope of another band-pass filtered white-Gaussian signal in the same frequency band as *y*(*t*). For instance, if *x*(*t*) is an alpha band oscillation and *n* = 2, *y*(*t*) is a beta band oscillation and 1:2 synchronized to *x*(*t*). If *a_x_*(*t*) = *a_y_*(*t*), the 1:n synchronization of these two signals computed from equation 2 is equal to 1. Note that in the case of *a_x_*(*t*) ≠ *a_y_*(*t*), the interaction of *x* and *y* is *for sure* genuine. Therefore, for the simulation of two genuinely (cross-frequency) synchronized sources, we used *a_x_*(*t*) ≠ *a_y_*(*t*).

The power of each oscillation is scaled based on the signal-to-noise (SNR) ratio of the frequency band of interest (see below).

##### Non-sinusoidal oscillations

A non-sinusoidal signal 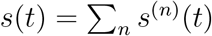, *n* ∈ *N* with the fundamental frequency of *f*_0_ was generated by adding up its fundamental component (or the first harmonic) *s*^(1)^(*t*) and the higher harmonics components *s*^(*n*)^(*t*), *n* ≥ 2. In the following equations, *s*^(1)^(*t*) is an oscillation at *f*_0_ produced by band-pass filtering a white Gaussian noise signal and *s*^(*n*)^(*t*), *n* ≥ 2 is a 1:n synchronized oscillation produced by equation 4 to be 1:n synchronized to *s*^(1)^.

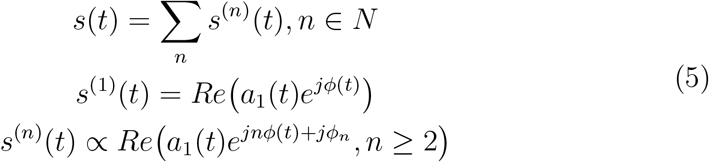

where *ϕ_n_, n* ≥ 2 are random numbers taken from a uniform distribution between [−*π/*2, *π/*2].

Given a fundamental frequency of *f*_0_, let 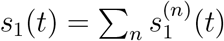 be a simulated non-sinusoidal oscillation based on equation 5 and 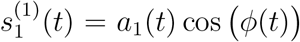. The following equations show how another non-sinusoidal signal *s*_2_(*t*) is simulated to be synchronized to *s*_1_(*t*):

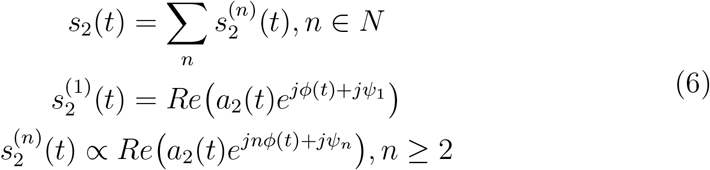

where *ψ_n_, n* ∈ *N* are random numbers taken from a uniform distribution between [−*π/*2, *π/*2]. In equation 6, 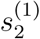 is an oscillation with 1:1 synchronization to 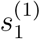

Note that the second harmonic is the strongest harmonic which is generally visible in real electrophysiological data. Therefore, without loss of generality, we only examine the removal of the second harmonic. Therefore, we simulated only the fundamental and the second harmonic. That is, in our simulations, the non-sinusoidal source signals are simulated as *s*(*t*) = *s*^(1)^(*t*) + *s*^(2)^(*t*) where *s*^(1)^(*t*) is an alpha oscillations and *s*^(2)^(*t*) is the second harmonic in beta frequency band. After that, the amplitude of *s*^(1)^(*t*) and *s*^(2)^(*t*) were re-scaled so that the SNR at each of alpha and beta frequency bands for these signals are set to the desired value (see below). Finally, *s*^(1)^(*t*) and *s*^(2)^(*t*) are added up together to generate *s*(*t*).

##### SNR

In realistic simulations, The SNR was defined as the ratio of the mean power of the source signal in the sensor space divided by the mean power of all pink noise sources in sensor space, filtered in the frequency band of interest. In our realistic simulations, the SNR of alpha and beta bands were set to 0dB and −10dB respectively.

For the toy examples, the SNR of a narrow-band source was defined as the ratio of its power to the power of the pink noise, filtered in the frequency band of interest. The SNR values at alpha and beta band were set to 5 dB and −5 dB respectively.

#### 2.5.2. Toy Examples

We used toy examples for initial assessment of the effect of Harmoni on the interactions between two signals with non-sinusoidal components. We used four scenarios for these toy examples, where the ground truth about the existing genuine and spurious interactions between the simulated signals were pre-defined. The left side of figure 5 depicts these scenarios schematically.

**Figure 5:**
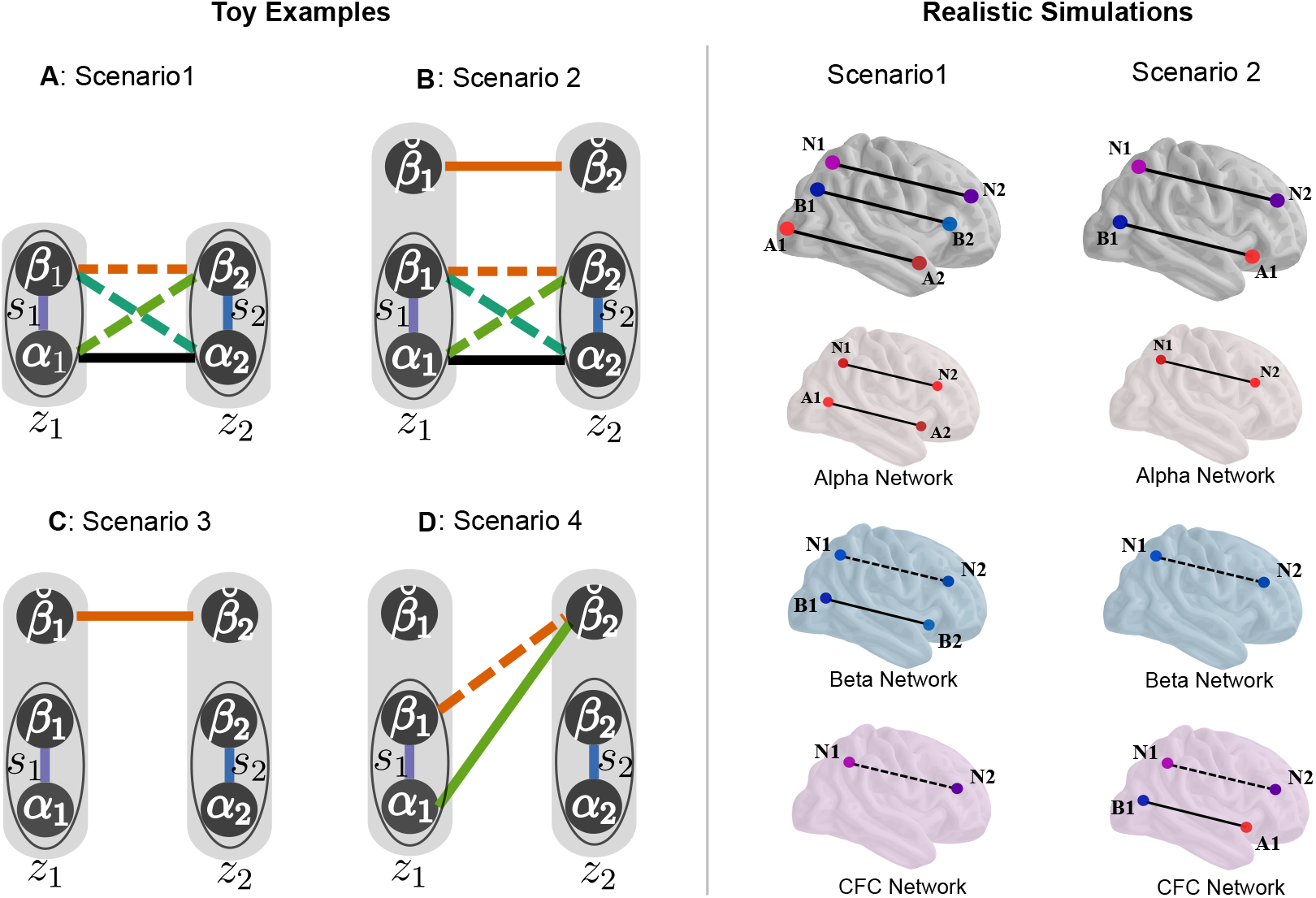
Simulation scenarios. Toy examples: Two signals *z*_1_ and *z*_2_ were simulated for each scenario, where various genuine and spurious synchronizations are present in the ground truth. The solid lines show the simulated, genuine synchronizations, and the dashed lines depict the spurious interactions observed in the ground-truth. Harmoni was applied on each of the signals and the within- and cross-frequency synchronization for alpha and beta bands were examined before and after Harmoni. In all scenarios, *z_k_* contained a non-sinusoidally shaped component *s_k_* = *α_k_* + *β_k_*, where *α_k_* and *β_k_* are the fundamental and second harmonic components of *s_k_* respectively. 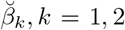 in scenarios 2 to 4 are beta oscillations independent of *s_k_, k* = 1, 2 Realistic simulations: In the first row, each dot shows a source and the connecting lines represent the synchronization of the source signals. The sources with purple color and the letter N correspond to sources with non-sinusoidal alpha oscillations having components in both alpha and beta frequency bands. The blue color and letter B corresponds to sinusoidal beta band sources, and the red color and letter A represent sinusoidal alpha frequency range sources. In the schematic brains of rows 2 to 4, the ground truth alpha, beta, and CFS networks are depicted. While solid lines depict genuine interactions, dashed lines show spurious interactions caused by non-sinusoidal waveshape of the signals. In both of the toy examples and realistic simulations, the main purpose of Harmoni is to suppress the spurious (dashed-line) connections, while not affecting the genuine (solid-line) interactions.

In each of the four scenarios, two signals *z_k_*(*t*), *k* = 1, 2 were simulated. On the schemes of figure 5, *z*_1_(*t*) and *z*_2_(*t*) are depicted as shaded areas in each scenario. In the rest of this section, the index *k* = 1, 2 refers to these two signals. *z*_1_(*t*) and *z*_2_(*t*) were multi-band signals with components in alpha and beta bands. In each scenario, specific ground truth genuine interactions were simulated between the two signals, which produced known spurious interactions, too. Harmoni was applied on each of the signals in order to remove the beta-component which could be the harmonic component of the alpha band component of the signal. The interactions between the two signals were estimated using absolute within- and cross-frequency coherence before and after Harmoni. We expected that Harmoni suppresses the spurious interactions, but does not touch the genuine interactions. For each scenario, 50 runs with random seeds were carried out.

In all scenarios, the two signals *z*_1_(*t*) and *z*_2_(*t*) contained an alpha oscillation with non-sinusoidal waveshape. *s_k_*(*t*) = *α_k_*(*t*) + *β_k_*(*t*) is the non-sinusoidal component of *z_k_*(*t*), where *α_k_*(*t*) represents the fundamental component and *β_k_* its second harmonic, which is phase-synchronized to *α_k_*(*t*).

Below, the composition of *z*_1_ and *z*_2_ in all the four scenarios and their genuine and spurious interactions are listed. Note that *ξ_k_*(*t*) is the additive 1/f (pink) noise component of *z_k_*(*t*).

##### Scenario 1 (figure 5-A)

*z_k_*(*t*) = *s_k_*(*t*) + *ξ_k_*(*t*), *k* = 1, 2. The signal *s*_1_ was simulated using equation 5 and *s*_2_ was simulated to be synchronized to *s*_1_ using equation 6. Therefore, a genuine interaction in alpha band between the two signals was simulated. Additionally, a spurious interaction in beta band, as well as spurious cross-frequency interactions between the two signals were observed in the ground truth. Figure 15 shows exemplar signals of this scenario.

##### Scenario 2 (figure 5-B)

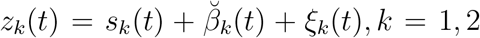. *s*_1_ and *s*_2_ were simulated as synchronized non-sinusoidal signals using equations 5 and 6 (similar to scenario 1). Each signal *z_k_* had an extra beta component 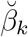. 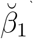 and 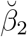 were simulated as narrow-band beta band oscillations and synchronized to each other (with equation 4) but independent of *s_k_, k* = 1, 2. In addition to the genuine integration between the *z*_1_ and *z*_2_ in beta band due to the synchronization of 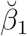 and 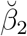, similar genuine and spurious interactions as in scenario 1 were present in the ground truth. In figure 15 15 an example of signals of this scenario is depicted (at the end of the manuscript).

##### Scenario 3 (figure 5-C)

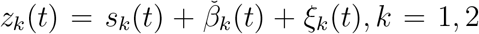. *s*_1_ and *s*_2_ were two independent non-sinusoidal oscillations (using equation 5) with their fundamental and second harmonic components in alpha and beta band respectively. 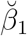 and 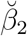 were two synchronized narrow-band beta oscillations (using equation 4), which were independent of *s*_1_ and *s*_2_. As a result, no CFS existed between *z*_1_ and *z*_2_ in the ground truth and the only genuine interaction was a synchronization within beta band.

##### Scenario 4 (figure 5-D)

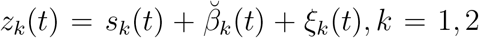. *s*_1_ and *s*_2_ were two non-sinusoidal alpha oscillations simulated independently using equation 5, and 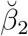 was a narrow-band beta oscillation 1:2 synchronized to *s*_1_, i.e. 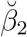 was simulated to have 1:2 CFS to the alpha component of *s*_1_ (*α*_1_) using equation 4. Therefore, in addition to the genuine CFS between *z*_1_ and *z*_2_, a spurious synchronization within beta band between *z*_1_ and *z*_2_ existed in the ground truth (i.e. between 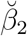 and *β*_1_). 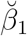 was a narrow-band beta oscillations independent of *s*_1_, *s*_2_, and 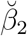.

Note that since there is no mixing between *z*_1_ and *z*_2_ in these simulations, the absolute coherence was used for quantifying both the within- and cross-frequency synchronizations.

#### 2.5.3. Realistic simulations

##### Source positions

The oscillatory sources were located at the center of randomly selected ROIs. Additionally, the position of 50 pink noise sources were selected randomly from the ∼8000 nodes of the source space grid. The Desikan Killiany (DK) atlas was used.

##### Scalp EEG generation

In order to generate the realistic multi-channel EEG signal, oscillatory and noise signals in source space were mapped to the sensor space using the forward solution with 64 electrodes according to BioSemi EEG cap layout. 200 datasets were simulated by using random seeds.

##### Realistic simulation scenarios

The two scenarios depicted on the right side of figure 5 were used for simulating realistic EEG data.

In scenario one, a pair of interacting non-sinusoidal source signals were simulated using equations 5 and 6 with their fundamental frequency in alpha band. Additionally, a pair of coupled sources in the beta band were generated using equation 4 and *n* = 1. A pair of synchronized sinusoidal sources in alpha band were simulated as well, by using equation 4 and *n* = 1.

In scenario 2, a pair of genuinely cross-frequency synchronized sources were simulated using equation 4 with *n* = 2. In addition, a pair of synchronized non-sinusoidal source signals were generated using equations 5 and 6.

##### Connectivity

The connectivity pipeline explained in detail above (also figure 4) was then applied to the simulated EEG data. As depicted in figure 5, each of these two scenarios include genuine and spurious interactions in their ground-truth. By using Harmoni, we expect to suppress the spurious interactions.

##### Evaluation criterion: ROC curve

Since the computed connectivity maps are not binary values (while the ground truth connectivity is binary), we evaluate the matching of computed connectivity maps and the ground truth using the area under curve (AUC) of the receiver operating characteristic (ROC) curve of the computed connectivity matrix. Figure 6 shows how true positive and false positive values are computed. After thresholding the test graph (*T*) with threshold level 0 ≤ *p* ≤ 1 (resulting in *T_p_*), The true positive ratio (TPR) and false positive ratio (FPR) corresponding to this threshold value are computed as 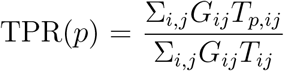 and 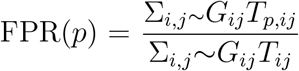, where the subscripts *ij* indicates the (*i, j*)-th element of the adjacency matrix and *G* is the ground-truth connectivity matrix. ∼*G* is the the 1’s complement of *G* (i.e., all zeros are converted to 1 and vice-versa).

**Figure 6:**
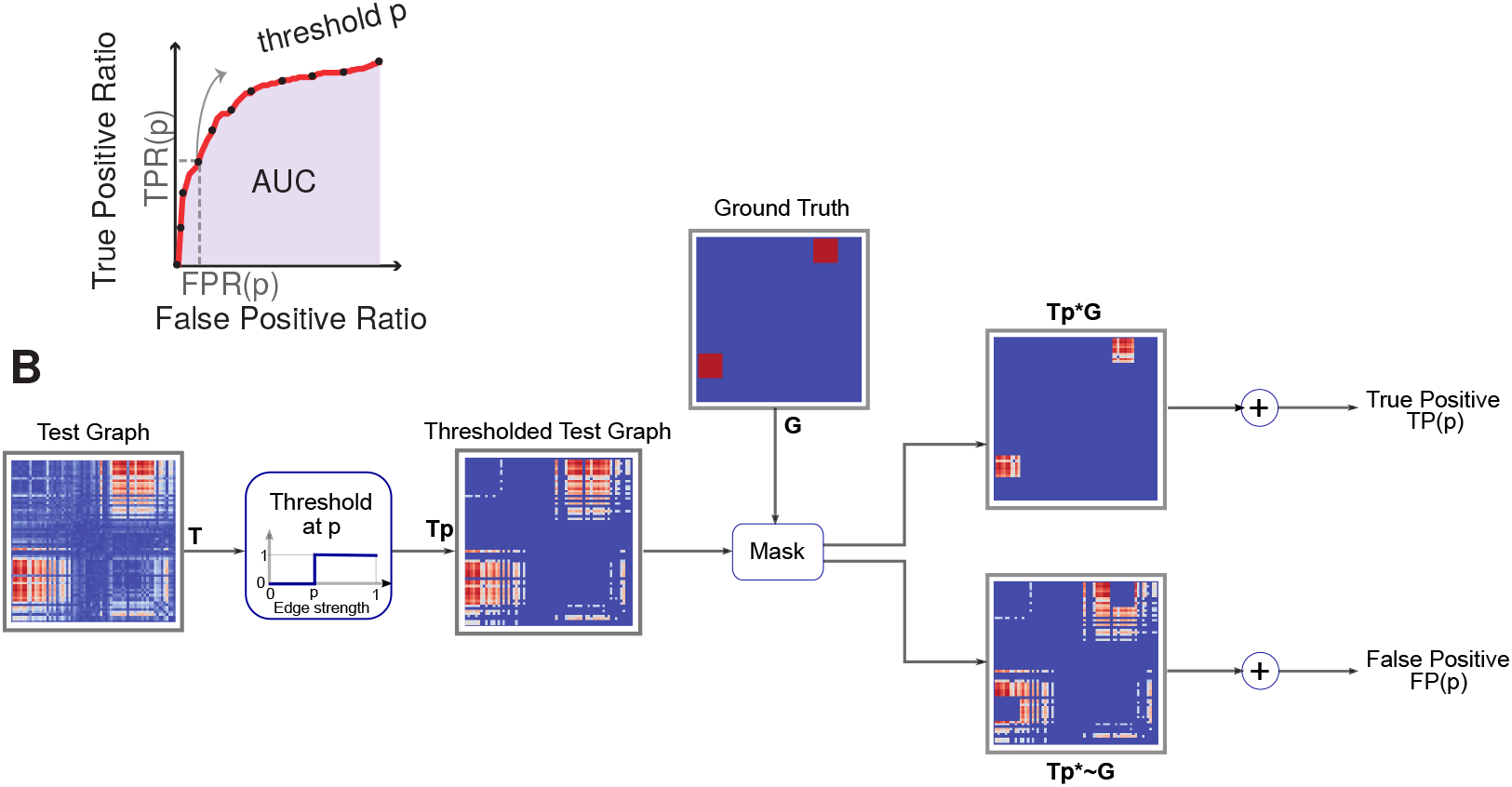
AUC of an ROC curve as an evaluation criterion for assessing the matching of computed connectivity graphs and the ground truth ones. Panel A shows an exemplar ROC curve. In panel b, the procedure of computing the true positive (TP) and false positive (FP) values corresponding to threshold level 0 ≤ *p* ≤ 1 is depicted. The true positive ratio (TPR) and false positive ratio (FPR) corresponding to each threshold level *p* is computed by 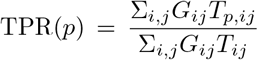 and 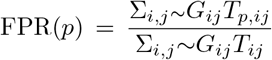. The *ij* index indicates the (*i, j*)-th element of the indexed matrix.

Using the TPR and FPR values for all the threshold level, an ROC curve is built. The AUC of this curve reflects how well the computed connectivity map matches the ground truth adjacency matrix of the graph corresponding to the simulated connectivity.

The AUC of the ROC curve (AUC-of-ROC) was computed for each simulation run before and after Harmoni and compared. We expected an increase of AUC-of-ROC after Harmoni.

Additionally, for graphs where no true positives were expected (for example the CFS network of scenario 1 or beta-band network of scenario 2) the FPR curve was built as a curve of FPR vs. threshold. The AUC of this curve (AUC-of-FPR) is a proxy of the amount of false positives. We expected a drop of AUC-of-FPR after Harmoni.

### 2.6. Resting-state EEG

#### 2.6.1. Data description

The resting-state EEG data from 81 subjects (20-35 years old, male, right-handed) of an open-access database (LEMON) were used (Babayan et al., 2019). The LEMON study was carried out in accordance with the Declaration of Helsinki and the study protocol was approved by the ethics committee at the medical faculty of the University of Leipzig. The data of each subject included 16 min resting-state recording with interleaved, 1-min blocks of eyes-closed and eyes-open conditions. For this manuscript, we used the data of the eyes-closed condition. The recordings were done with a band-pass filter between 0.015 Hz and 1 kHz and a sampling rate of 2500 Hz.

For our analysis, we used the publicly available preprocessed data in the database. The sampling rate was reduced to 250 Hz and the down-sampled data were filtered within [1, 45] Hz with a fourth order Butterworth filter, applied forward and backward. Then the data segments of eyes-open and eyes-closed conditions were separated. Bad segments were removed manually and ICA artifact rejection was employed to remove the noise components relating to eye, heart, and muscle activity. Babayan et al. (2019) provide detailed information about the data recording and preprocessing steps.

#### 2.6.2. Connectivity

The pipeline in figure 4 was used, as simular to the simulated data connectivity. Fourth-order Butterworth filters (applied forward-backward to avoid phase shift) were used for filtering data in alpha band (8-12 Hz) and beta band (16-24 Hz). Similar to the connectivity pipeline described in detail above (also figure 4), the band-pass filtered data were then projected onto cortical source space using the inverse solution computed from fsaverage standard head, with 4098 vertices per hemisphere. Afterwards, a single time series was extracted (using SVD) for each ROI from the cortical sources within that ROI. The Schaefer atlas (Schaefer et al., 2018) with 100 ROI and 7 Yeo resting-state networks (Yeo et al., 2011) was used.

For each subject, the ROI-ROI connectivity for alpha-beta CFS was computed before and after Harmoni, resulting in 100 × 100 connectivity adjacency matrices. In order to make the connectivity graphs comparable before and after Harmoni at the group level, the adjacency matrix of each subject was z-scored before and after Harmoni. The z-scored matrices of the networks before Harmoni were subtracted from the ones after Harmoni. Two-sided paired t-tests was used for each connection to specify the links which were changing significantly on group level. The Bonferroni method was used to correct for multiple comparisons, i.e. the p-values were multiplied by 100^2^ and then the links with corrected pvalues > 0.05 were considered as significant.

##### Asymmetry-index of CFS networks

In order to quantify the extent to which the CFS adjacency matrices are asymmetric, we used the norm of the anti-symmetric part of the adjacency matrix. For a given matrix **A**, the antisymmetric part is defined as 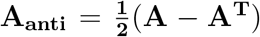. We define 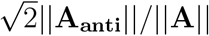 as an *asymmetry-index*. It can be shown that this index is between zero and one, with zero value corresponding to a symmetric matrix and a value of one for an antisymmetric matrix.

### 2.7. Depiction of CFS connectivity

We used a bipartite graph for the depiction of CFS networks. The CFS networks have an asymmetric adjacency matrix and therefore, should be depicted as directed graphs. We actually used a bipartite graph as a way of illustrating a directed graph in a more comprehensive way.

A bipartite graph is a graph which has two sets of nodes and an edge can only connect the vertices from different sets (i.e. alpha and beta sets in our analysis) to each other. In our case of CFS networks, each node is a representative of a brain region and each set of nodes relates to the activity of the brain regions in one of the frequency bands. Figure 7 shows an illustrative example of such depiction for alpha-beta CFS. The upper and lower node-sets represent the alpha and beta band activity of the ROIs of interest, respectively. A link between node 1 from the upper set (alpha nodes) with node 3 of the lower set (beta nodes) shows a CFS coupling between ROI 1 and 3. This connection would be the element (1,3) of the adjacency matrix of the network. In a directed graph this edge would be an out-going edge for node 1 and an in-coming edge for node 3.

**Figure 7:**
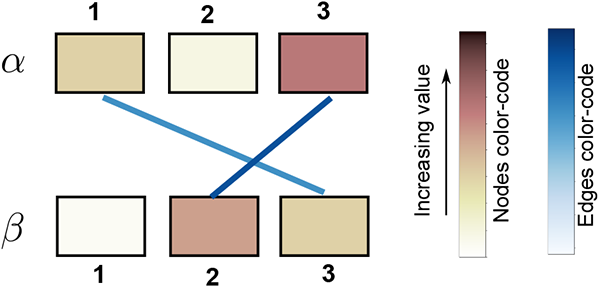
Depicting CFS network as a bipartite graph. The nodes stand for brain regions. While the upper set of nodes represents the alpha activity in the brain regions, the lower nodes are for the beta activity in those regions. When node 1 from alpha nodes (upper nodes) is connected to node 3 of beta nodes (lower nodes) it means that the alpha activity in region 1 is coupled to beta activity in node 3. The links are color-coded based on the strength of the coupling. Additionally, each node in each frequency band can have a color which represents its centrality in that frequency band.

In our illustration of the graph, each node can have a color, which shows its centrality value. In this work, we did not use this feature and the node colors are the label colors provided with the parcellation. For real data these colors code the ROI’s Yeo resting-state network. Each edge is also color-coded with the strength of the coupling that it represents. It can be the absolute or relative strength of coupling.

### 2.8. Statistical Analysis

Two-sided paired t-tests were used for testing the difference of the mean value of two paired samples. Specifically, the changes of the evaluation parameters in simulations (the AUC values) as well as real data (the change in the connectivity values and the asymmetry-index) were tested before and after Harmoni.

For testing the significance of the correlation of the initial value of a parameter (before Harmoni) and its percentage change after Harmoni, we used the correction method introduced in (Tu, 2016). Assume *x* is the baseline value of a parameter of interest before Harmoni and *y* is its value after Harmoni. The percentage change of this parameter is defined as (*y* − *x*)*/x*, which is mathematically coupled to *x*. Therefore, it would not be valid to use the conventional statistical testing between the initial value and the percentage change and compare the observed correlation to zero. Tu (2016) suggests that the appropriate null value for the hypothesis test should be 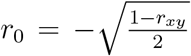 rather than zero, where *r_xy_* is the Pearson correlation of *x* and *y*. In this approach, the hypothesis test is 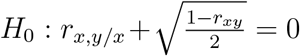 versus 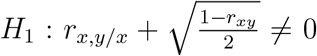. Finally, the expression for the z-test is suggested to be 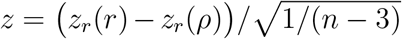, where *z_r_*(*r*) = 0.5*ln*((1 + *r*)/(1 − *r*)) is the Fischer’s z transformation, *r* is the observed correlation coefficient, and *ρ* is the correlation coefficient to be tested against.

## 3. Results

### 3.1. Simulations

#### Toy Examples

As the very first step, we used simplified simulations (toy signals) to show that Harmoni is an effective algorithm for suppressing spurious CFS and within-frequency interactions due to the non-sinusoidal shape of the signals. In these simple simulations, where there are no complications regarding source mixing or limitations of source reconstruction, the ground truth about the interactions between the two simulated signals is known. In fact, we were interested to validate two important properties of Harmoni: (1) It suppresses the spurious interactions significantly, and (2) it does not affect genuine interactions. In addition, these initial simulations serve as a demonstration for the main spurious interactions present due to non-sinusoidality.

In each of the four scenarios, two noisy multi-band signals *z_k_*(*t*), *k* = 1, 2 were simulated with components in alpha and beta band. Different genuine interactions were simulated between the two signals, resulting in spurious interactions as well. Harmoni was applied to each of the two signals to remove beta components associated with being a harmonic of alpha band components, i.e. showing CFS with the alpha oscillation. The within- and cross-frequency interactions were then estimated using absolute coherence to investigate how they changed after using Harmoni and how these changes were related to the ground truth. Each scenario was simulated 50 times with random seeds. Figure 8 depicts the boxplots of the strength of possible within- and cross-frequency interactions between and within the two signals, before and after Harmoni. The interactions in the schematic of each scenario have the same color-code as their respective boxplots. The change of the synchronization strength after Harmoni (in comparison to before Harmoni) was tested with a two-sided paired t-test for each possible interaction, and then corrected by the Bonferroni method.

**Figure 8:**
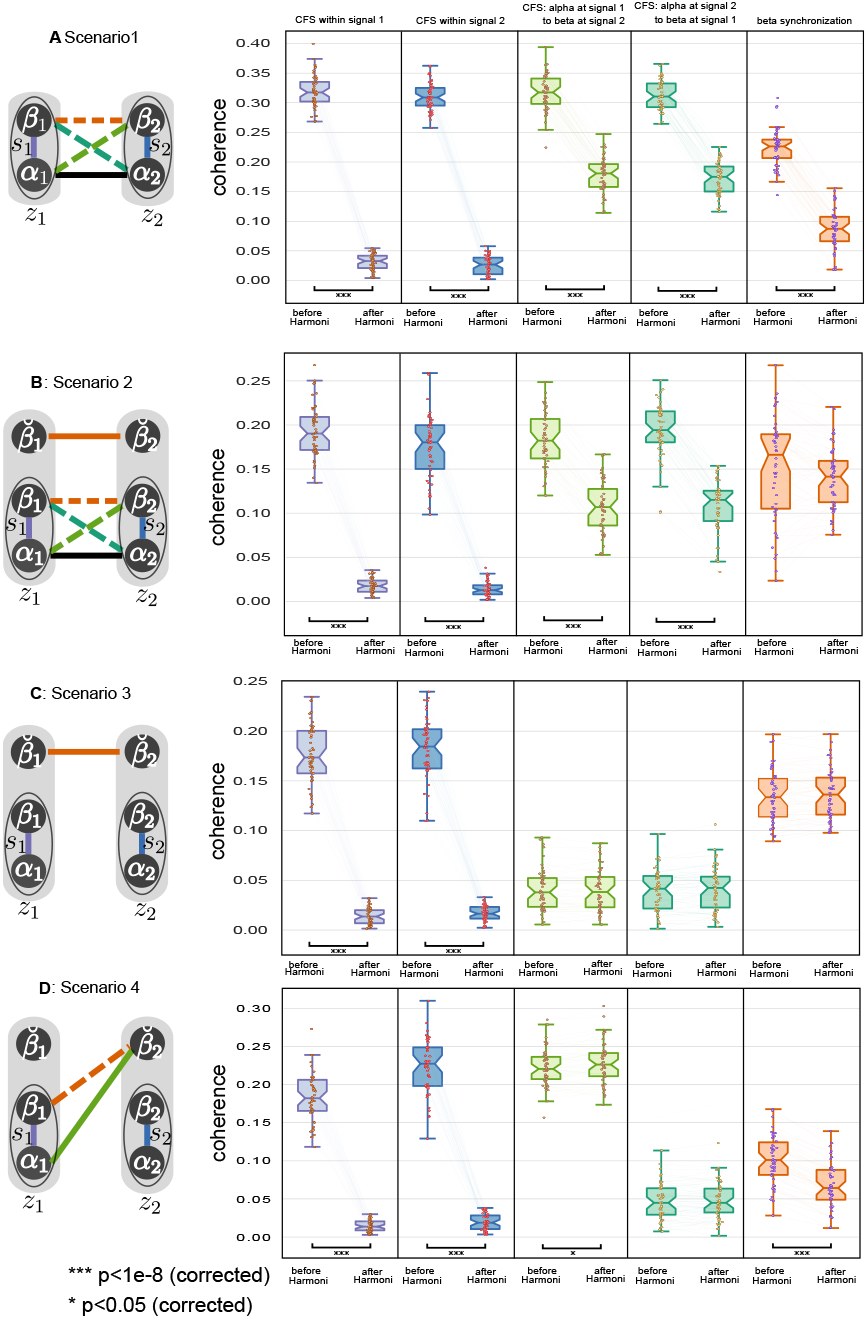
Performance of Harmoni on toy examples in 50 runs with random start seeds. The left -side schemes are the simulation scenarios shown in figure 5. For all scenarios the strength of each possible interaction is shown before and after Harmoni in the boxplots in the same panel as the scenario scheme. The purple and blue color are associated with the within-signal CFS, the two green colors are related to the inter-signal CFS values, and finally the orange color is dedicated for the beta band synchronization among the two signals. In all scenarios, two signals are simulated and each of them contains a non-sinusoidal wave *s_k_*(*t*) = *α_k_*(*t*) + *β_k_*(*t*), *k* = 1, 2 with their fundamental component *α_k_* in alpha band and their second harmonic *β_k_* in beta band. Scenario one: The boxplots show that all of the within-signal CFS and the spurious interactions are suppressed significantly. Scenario two: Only the beta-synchronization between the two signals does not change significantly after Harmoni and stays at a large value due to the genuine synchronization of 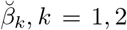. Scenario three: The CFS within each signal is suppressed significantly, the CFS values between the two signals do not change and have small values in general, and importantly the beta-synchronization between the two signals stays almost the same at a high value. Scenario four: a genuine CFS (light green) between the two signals is simulated, which is not affected after Harmoni, while the spurious within-beta interactions and the within-signal CFS are suppressed.

In scenario one (figure 8-A), the two signals were synchronized non-sinusoidal waves with their fundamental frequency in alpha band (i.e., *z_i_*(*t*) ≈ *s_k_*(*t*) + *ξ_k_*(*t*) with *s_k_*(*t*) = *α_k_*(*t*) + *β_k_*(*t*) being the non-sinusoidal component of *z_k_*(*t*). *s*_1_ and *s*_2_ were simulated to be synchronzied, i.e. *α*_1_ ⟷ *α*_2_, where α shows the synchronization). The CFS interaction between the two signals as well as the interaction in beta band are by construction spurious. As shown in figure 8-A, the within- and cross-frequency spurious coherence between and within the two signals are successfully suppressed after Harmoni.

In scenario two (figure 8-B), each of the two signals contained another beta component which was independent of the non-sinusoidal components, but these components from *z*_1_ and *z*_2_ were simulated to be synchronized to each other (i.e., 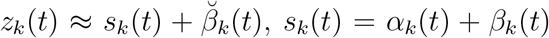, with *α*_1_ ⟷ *α*_2_, 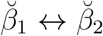). In this scenario, the CFS interaction is by construction spurious, too. However, a part of the interaction between the two signals within the beta band is genuine because of the interaction between 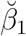 and 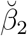. The results in figure 8-B show that the CFS interactions are suppressed, and the coherence between the beta components of the two signals does not have any significant change, showing that the genuine beta synchronization is still present..

Scenario three (figure 8-C) was similar to scenario two with the difference that the non-sinusoidal oscillations from the two signals were not synchronized (i.e., 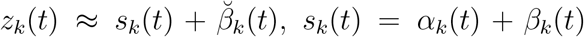, with 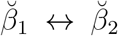). Therefore, no CFS between the two signals is observed. The boxplots in figure 8-C show that the CFS within each signal is suppressed as expected from the proper functioning of Harmoni, while CFS between the two signals does not change, remaining at a negligible level. Importantly, the genuine synchronization in beta-band does not change after Harmoni.

In scenario four (figure 8-D) 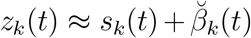, *s_k_*(*t*) = *α_k_*(*t*) + *β_k_*(*t*) as well. The ground truth interactions were set to 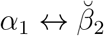. This setting results in genuine CFS between the two signals. Figure 8-D shows that Harmoni is robust: the genuine inter-signal CFS does not change, while the present CFS within each signal as well as the spurious beta-band interaction drop significantly. Additionally, the other CFS between the two signals which was missing by construction, does not change and remains at a low value.

All in all the results of the above scenarios show that the spurious interactions are suppressed by Harmoni, while the genuine interactions are not changed.

##### Realistic EEG simulations

For the further evaluation of Harmoni, we developed an EEG simulation pipeline for generating realistic scalp EEG signals (details in the method section). The simulated EEG data consisted of narrow-band sinusoidal source signals at alpha (8-12 Hz) and beta (16-24 Hz) bands, as well as non-sinusoidal signals with fundamental frequency at alpha band. The dipole positions were randomly selected from the center of 68 regions of interest (ROIs) of Desikan Killiany atlas (Desikan et al., 2006). 1/f (pink) noise data were also added to the generated source signals of interest. All the source signals were forward modelled to generate realistic EEG. Two scenarios (shown in figure 5) were used for generating the simulated EEG signals. Both of the scenarios included coupled non-sinusoidal alpha sources. In scenario one there were also within-frequency coupled narrow-band sinusoidal alpha and beta sources. In scenario two, in addition to the pair of coupled non-sinusoidal sources, a genuine, remote cross-frequency coupled pair of sinusoidal sources was simulated as well. As shown in figure 5, these two scenarios have differential within- and cross-frequency network profiles.

We used the connectivity pipeline of figure 4 to compute the within-frequency synchronization in beta band and the alpha-beta cross-frequency synchronization maps.

As an illustrative example (figure 9) and a proof of principle, we first show an example of scenario two. Two synchronized non-sinusoidal alpha source signals were simulated with their corresponding sources in caudal middle-frontal and inferior-parietal regions of right and left hemispheres, respectively. In addition, two sinusoidal alpha and beta source signals, with CFS, were simulated in the caudal middle-frontal and inferior-parietal regions of the left and right hemispheres, respectively. The ground truth networks are shown in figure 9-A. Afterwards, the source signals, along with random noise sources, were projected to the sensor space and then the above-mentioned source space pipeline was performed. Panel B of figure 9 depicts the top 1% connections of the connectivity networks in alpha band as well as beta band and CFS networks before and after Harmoni. The spurious beta and CFS connections are suppressed.

**Figure 9:**
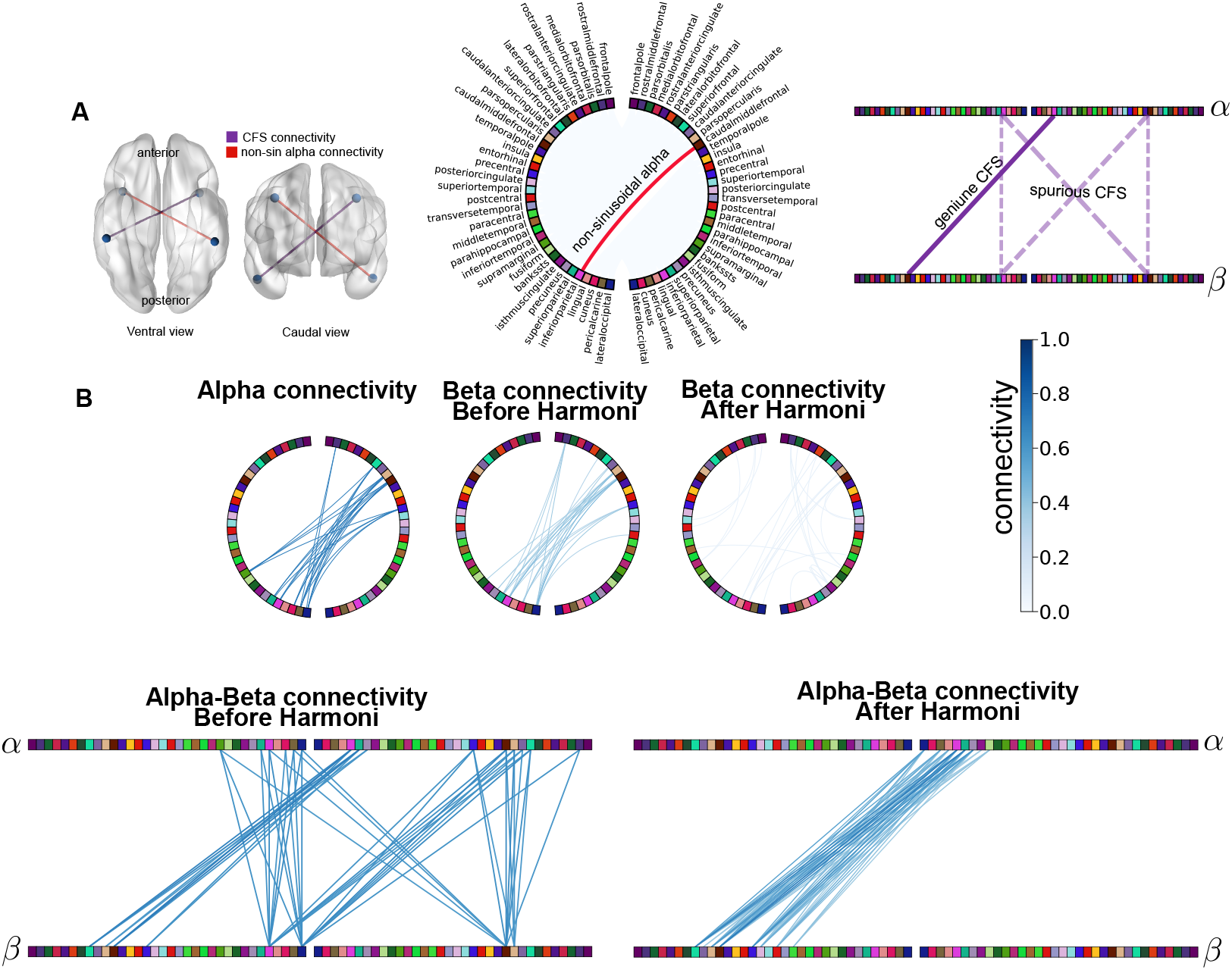
An illustrative realistic simulation example, showing the effect of Harmoni in suppressing the spurious interactions due to harmonics. Panel A depicts the ground truth, where synchronized non-sinusoidal alpha sources were simulated in right caudal middle-frontal and left inferior-parietal regions (red connecting line) and two cross-frequency synchronized narrow-band alpha and beta sources were simulated in the left caudal middle-frontal and right inferior-parietal regions (purple connection). The circular and bipartite graphs depict the ground truth alpha and CFS networks. A bipartite graph allows to see how different nodes from two networks, represented by horizontal bars, connect to each other allowing non-symmetric connections - without using a directed graph. In the CFS network, the dashed-lines represent the spurious interactions due the connectivity between two non-sinusoidal signals, while the solid line represents the genuine interaction. Panel B shows the top 1% connections of the within-frequency and cross-frequency networks computed before and after Harmoni. The spurious beta connections and the spurious CFS connections are suppressed. The glass brains were plotted with Brain Network viewer (Xia et al., 2013) in MATLAB. The circular plots were generated with MNE Python (Gramfort et al., 2013, 2014)

Our main evaluation criterion for the realistic simulations was the area under curve (AUC) of the receiver operating characteristic (ROC) curve and the false positive ratio (FPR) curve. These curves were built by comparing the adjacency matrix of the connectivity graphs before and after Harmoni to their counterpart ground truth connectivity matrices. The ROC curve was computed for the beta network in scenario one and the CFS network in scenario two. The higher the AUC of ROC curve (AUC-of-ROC), the more similar the connectivity matrix to the ground truth one. Figure 10 shows the results of evaluating the two scenarios of the simulation in 200 Monte Carlo simulations with random dipole positions. The increase of the AUC-of-ROC in the left sides of panels A and B demonstrates a success of Harmoni in both of the scenarios in correcting the connectivity maps in the way that they are more similar to the ground truth. Consequently the ratio of the true positive ratio (TPR) and FPR increases after Harmoni, reflecting the suppression of spurious interactions (false positives) and not affecting/increasing the genuine interactions (true positives). Moreover, the percentage change of the AUC-of-ROC values decreases with the increase of the initial value of AUC-of-ROC. That is, the closer the initial connectivity map to the ground truth, the less correction Harmoni applies. In other words, if a network shows a lot of spurious interactions, then it is corrected by Harmoni more strongly (see statistical analysis section in Methods for quantifying this dependency in a statistically stringent manner). In addition, at the left sides of both the panels of figure 10 the AUC of the FPR curves (AUC-of-FPR) of the CF networks in scenario one, and the beta networks in scenario two (where all the present interactions are spurious) decrease after Harmoni (the second columns in figure 10-A and B), showing the suppression of the spurious interactions. The absolute value of the percentage change of the AUC-of-FPR in these cases increases with the increase of the initial value. This means that the more false positive links are present in the connectivity maps, the more pronounced is the impact of Harmoni on the networks.

**Figure 10:**
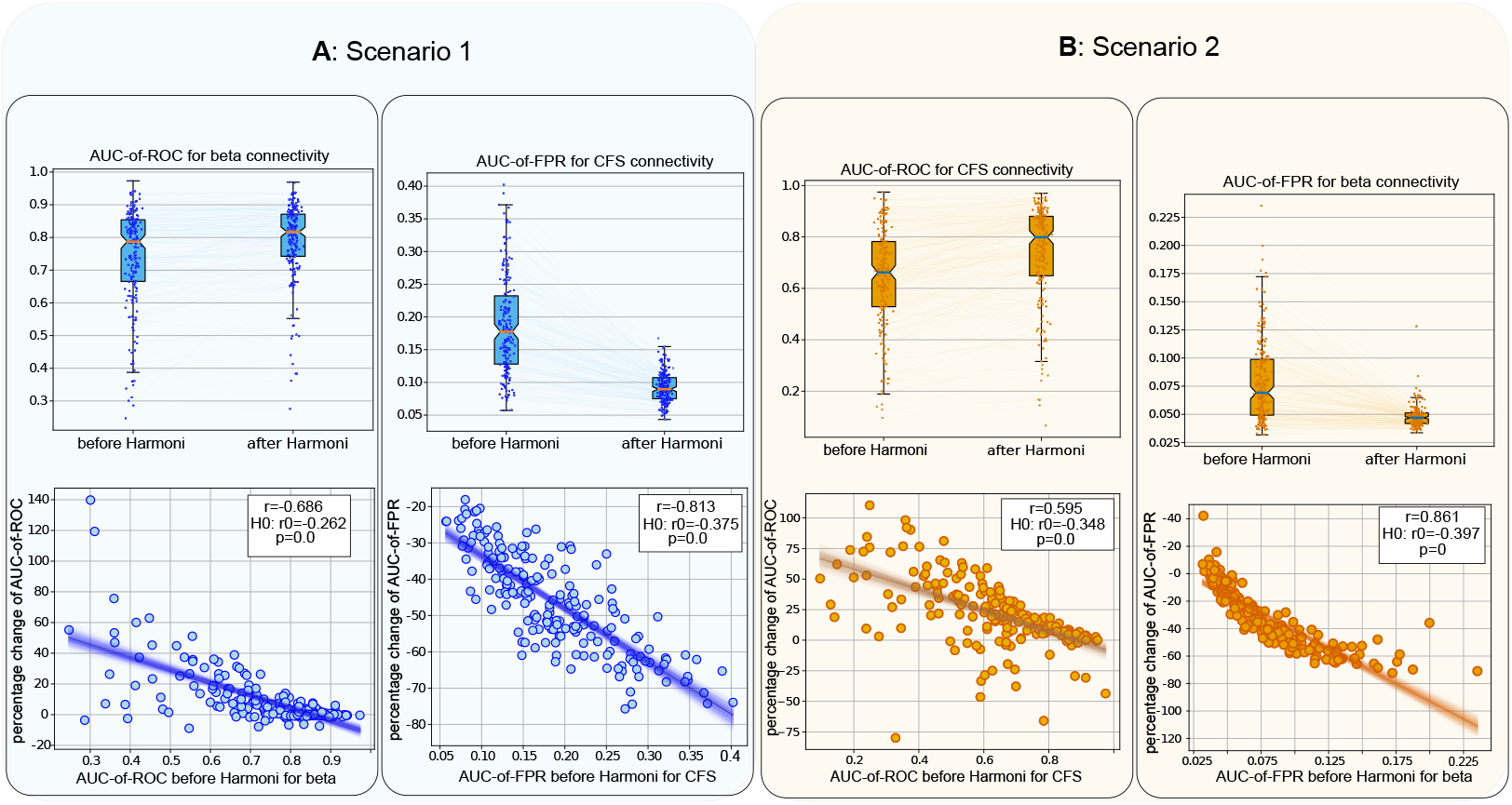
Results of 200 realistic simulations according to scenario one (panel A) and two (panel B) of figure 5. At the left side of panel A, the boxplots of the AUC-of-ROC of beta connectivity before and after Harmoni are depicted, showing an increase after the application of Harmoni. This indicates a successful correction of the network’s connections after Harmoni in favor of suppressing the spurious interactions. Beneath the boxplots, the scatter-plot of the percentage change vs. the AUC-of-ROC values for beta connectivity before Harmoni is shown. The higher the initial AUC-of-ROC value (i.e. the more accurate the initial connectivity map), the less difference between the AUC values before and after Harmoni (i.e., the less the impact of Harmoni). At the right side of panel A the boxplots of the AUC-of-FPR for the CFS connectivity are illustrated. Note that in scenario one the whole CFS connectivity is spurious due to waveshape, which is to a great extent removed by Harmoni (reflected in the decrease of the FPR). The bottom scatter-plot shows that the percentage change increases as the AUC-of-FPR of the CFS network increases, meaning that Harmoni has a larger effect on networks with more spurious interactions. Panel B shows the results of scenario two, but for the AUC-of-ROC of the CFS network (the left side) and the AUC-of-FPR of the beta connectivity (the right side). A similar outcome as in scenario one is observed in scenario two: and increase in the AUC-of-ROC after Harmoni for CFS networks, as well as a decrease in AUC-of-FPR for beta networks where all the connections are spurious ones. The percentage-change scatter plots imply a similar effect: the more spurious interactions in the simulated signals, the more corrections is performed by Harmoni.

### 3.2. Harmoni on resting-state EEG data

Alpha oscillations recorded with resting-state EEG (rsEEG) are known to have a non-sinusoidal waveshape in many brain areas. For example, the *μ* rhythm in the somatomotor areas or visual alpha are well-known examples of non-sinusoidal oscillations. This non-sinusoidal waveform is manifested in the power spectral density (PSD) having a large peak at alpha and a smaller peak at beta frequency band, together with 1:2 CFS between alpha and beta bands. As an example from real data, figure 11 shows a segment of a non-sinusoidal source signal extracted from the recordings of a subject’s eyes-closed rsEEG from the LEMON dataset (Babayan et al., 2019). In this case, the power spectrum of such signal shows two prominent peaks at the fundamental frequency (11Hz) and its second harmonic frequency (22Hz). Additionally, a third peak is visible at the third harmonic frequency as well (33Hz). As indicated by the values of the cross-frequency coherence in the figure, the harmonic components demonstrate CFS with the fundamental frequency component.

**Figure 11:**
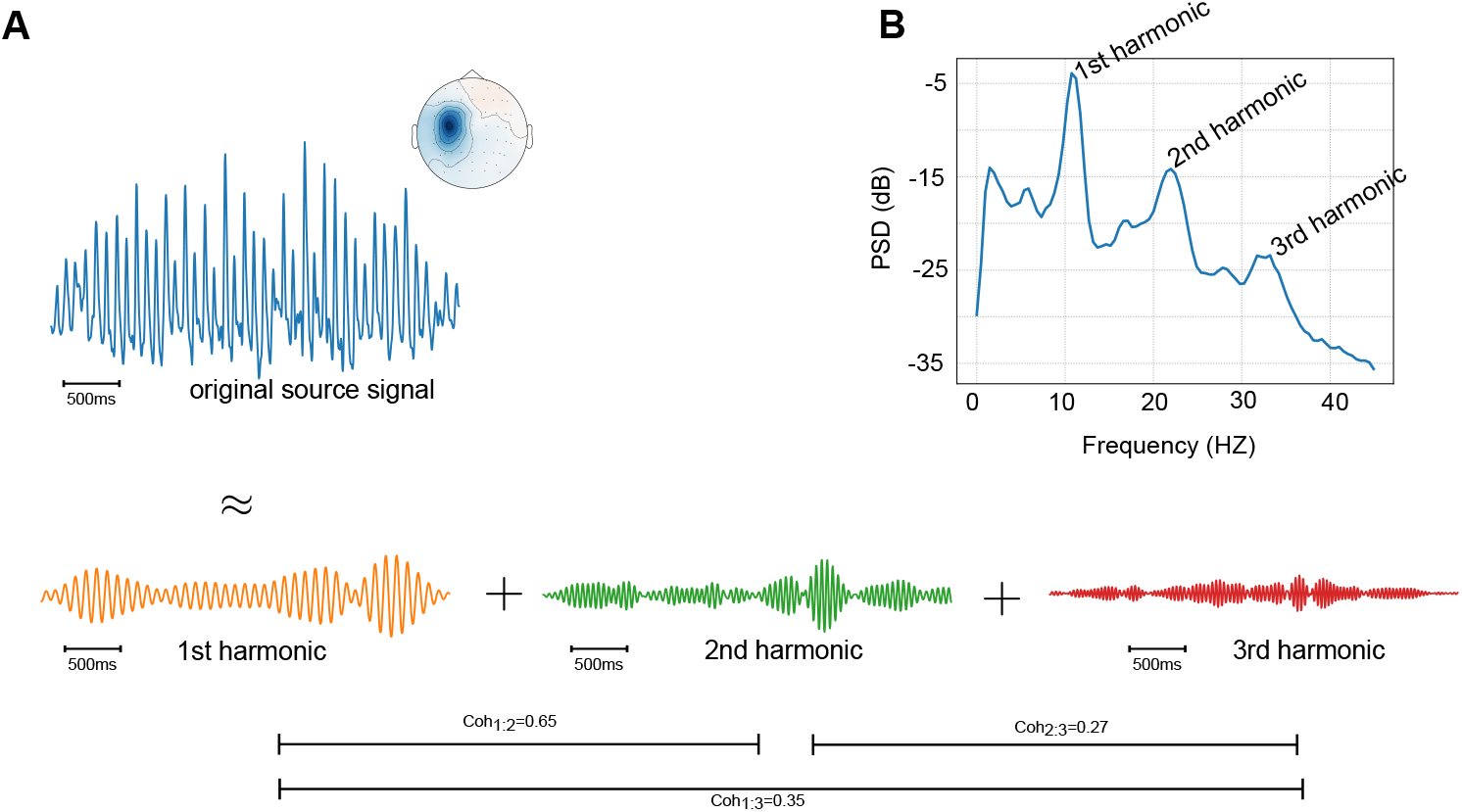
An example of a non-sinusoidal brain source signal. In panel A, a non-sinusoidal brain oscillatory activity and its first three harmonics are shown along with the spatial pattern of this activity. This source was extracted from eyes-closed rsEEG of a subject of the LEMON dataset using independent component analysis (ICA) (extended InfoMax ICA (Lee et al., 1999) with 32 components). Panel B shows the PSD of the non-sinusoidal signal with the peaks at 11 Hz (first harmonic, or the fundamental frequency), 22 Hz (second harmonic), and 33Hz (third harmonic). The cross-frequency coherence of the harmonic components and the fundamental component are reported as well. The largest synchronization occurs between the first and second harmonic (coherence value of 0.65). This is mainly due to the higher signal-to-noise ratio at these frequency bands.

We used rsEEG data from 81 subjects (data description in the Method section) and applied Harmoni in order to disambiguate genuine from spurious CFS alpha-beta interactions. Panel (A) of figure 12 illustrates the across-subjects average of 1:2 alpha-beta synchronization at each cortical source (i.e. a vertex on the cortical mantel). A very high 1:2 synchronization within one cortical source is an indication of a non-sinusoidal waveshape of alpha oscillation at the corresponding dipole. On average, the occipital, temporal and central areas demonstrate the highest 1:2 alpha-beta synchronization. This figure shows the ubiquity of harmonics in data and highlights the importance of taking care of it in connectivity analysis. Note that although we make the assumption that the 1:2 synchronization at a single source is a harmonic-driven synchronization, we are fully aware that this can be a result of residuals of signal mixing in source space. We explicitly address this point in the discussion.

**Figure 12:**
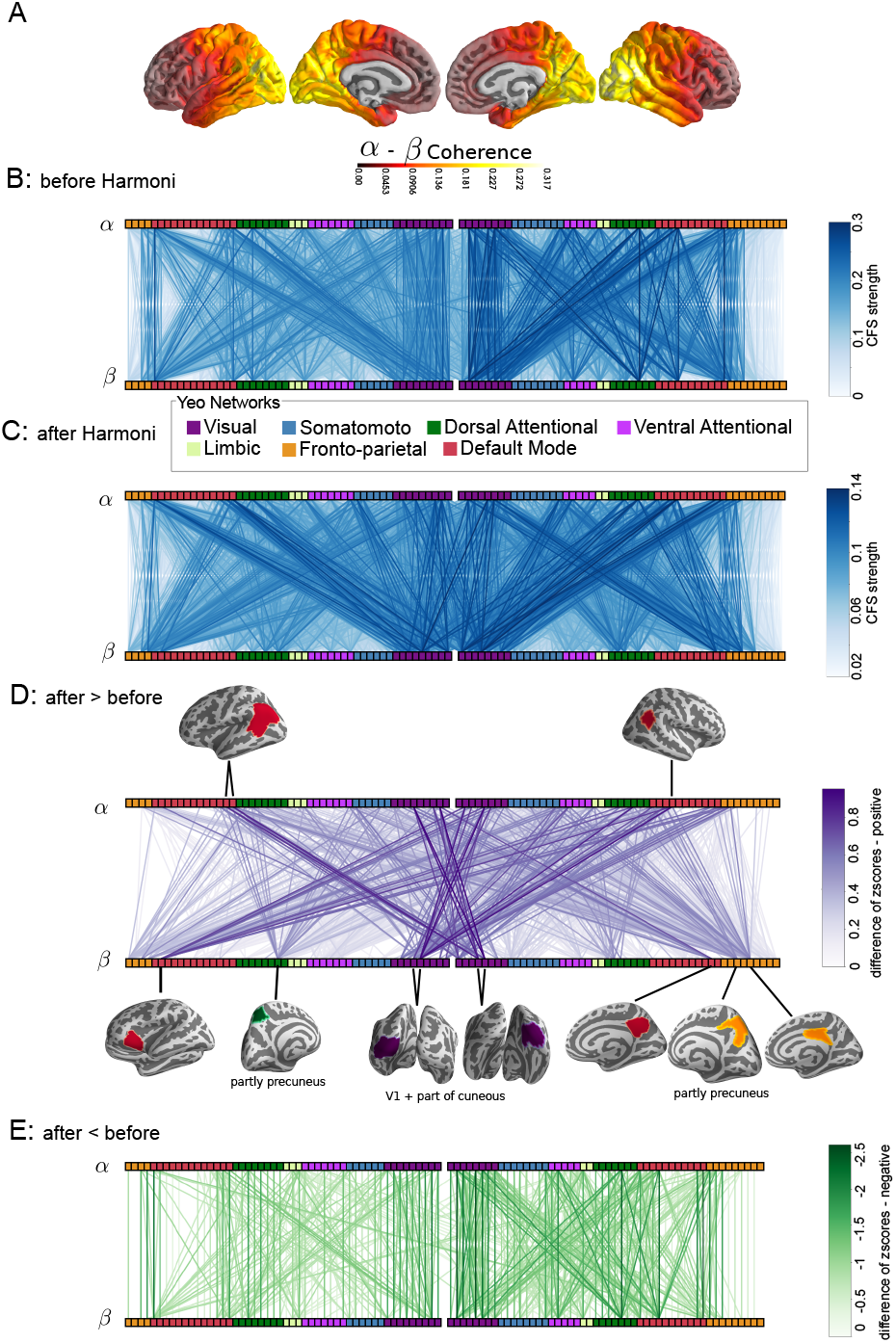
Harmoni and rsEEG data. Panel (A) shows the across-subject average of 1:2 synchronization of the alpha and beta band activity over the cortex. If the 1:2 synchronization is high at a given source, the second harmonic of the alpha activity may have a large contribution to the beta activity. Panel (B) shows the bipartite illustration of the mean CFS connectivity matrix. The nodes are sorted based on their assigned Yeo resting-state network. The vertical links show the presence of CFS within a single region, which is a sign of a synchronization due to waveshape (since this way they connect the same region). Panel (C) is similar to panel (B), but for the data after the application of Harmoni on beta band. The vertical links in the bipartite illustration are eliminated and more inter-hemispheric connections emerged. Panel (D) shows the links which are more pronounced after Harmoni, including more inter-hemispheric interactions. Panel (E) shows the links which were suppressed by Harmoni. The networks of panels (D) and (E) were computed by subtracting the z-scored coherence values before Harmoni from the ones after Harmoni.

In order to compute the CFS connectivity networks, a similar data-analysis pipeline as in the realistic simulations was used at the source space. The rsEEG multi-channel data were mapped to 100 ROIs of the Schaefer atlas (Schaefer et al., 2018) with each ROI being assigned to one of the seven resting-state Yeo networks, i.e. Default-mode network, Fronto-parietal, Limbic, Ventral Attentional, Dorsal Attentional, Somatomotor, and Visual networks (Yeo et al., 2011). Then, the components of beta activity at each ROI that could potentially be a higher harmonic of alpha oscillations were removed using Harmoni. Finally, the ROI-ROI alpha-beta CFS connectivity networks, represented by 100 × 100 connectivity matrices were computed. Figure 12-B and C show the across-subject mean connectivity graphs before and after Harmoni over all subjects. In Panel B (CFS before Harmoni), the dominating vertical links correspond to the local synchronization of the alpha oscillations with their second harmonic (beta). This is an expected pattern for the non-sinusoidal oscillations where both alpha and beta components are generated at the same location and demonstrate spurious CFS. Panel C shows that the application of Harmoni resulted in the unmasking of genuine remote neuronal interactions which were previously under-emphasized due to the presence of spurious cross-frequency connectivity. In order to be able to compare the networks before and after Harmoni at the group level, the connectivity matrices were z-scored for each subject and then these standardized coherence scores before Harmoni were subtracted from the ones after Harmoni, and paired two-sided t-tests (with Bonferroni correction of p-values) were employed to specify the links which changed significantly after Harmoni. Panel 12-D and E show the across-subject mean of the difference networks for positive and negative links (only the significantly changing links). 12-D depicts the connections which are more pronounced after Harmoni. This enhancement is observed for both inter and intra-hemispheric connections, specifically between the visual cortices of the two hemispheres, between the visual areas and the default mode and fronto-parietal regions. These effects were achieved via the elimination of spurious connections which were driven by harmonics. The presence of such harmonics masks the strength of the genuine interactions which, however, become more pronounced after the application of Harmoni. The presence of vertical lines and some cross-region lines in figure 12-E illustrates that within-ROI CFS as well as many within-hemispheric connections are significantly suppressed.

Importantly, Harmoni does not create any new connections, it rather leads to a reweighing of the connections after the suppression of the spurious ones. In order to validate this claim, we used paired t-tests to check whether the across-subject mean of the weights of each connectivity link changes significantly after Harmoni. Accounting for multiple comparisons by Bonferroni correction, we found that all the significant changes were in the direction of a decrease in the connectivity strength after Harmoni, −12.77 ≤ *t*(80) ≤ −4.9, *p* < 0.05 (figure 13), which confirms that no *new* connection is produced by Harmoni. Indeed, by suppressing the synchronizations that can mimic the spurious interactions due to non-sinusoidal waveshape of alpha oscillations, the ratio of the connectivity weights with respect to the maximum synchronization is changed and therefore, some connection weights which previously were in the low ranks move to higher percentiles of the connectivity weights after the application of Harmoni. With this procedure, the dominant and strongest connections change in the CFS network and we observe the networks in figure 12-B and C.

**Figure 13:**
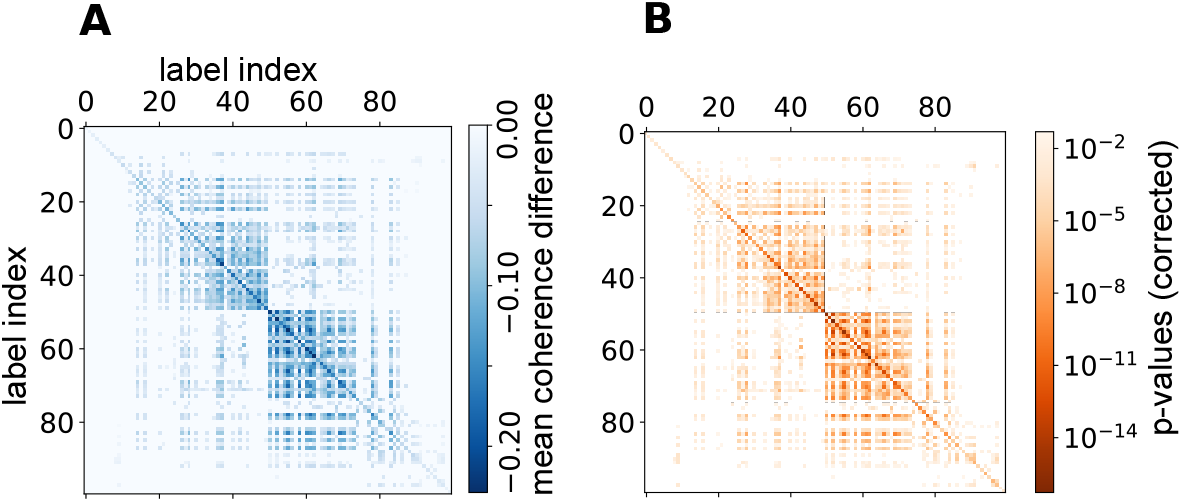
Harmoni does not create new connections, i.e., an appearance of a synchronization between two ROIs after Harmoni which was not present before Harmoni. Panel (A) shows the significant across-subjects mean difference of the alpha-beta networks after and before Harmoni (the coherence values before Harmoni were subtracted from the values after Harmoni). All the values are ≤ 0, showing that the synchronization strengths drop for all pairs of the ROIs on average. (B) The matrix of corrected p-values (Bonferroni corrected) corresponding to the two-sided paired t-tests performed for each CFS connection before and after Harmoni. The insignificant connections are not colored. All the significant changes indicated a decrease, − 12.77 ≤ *t*(80) ≤ −4.9, *p* < 0.05 (after Bonferroni correction).

Another important feature of the MEG/EEG connectivity networks is the symmetry of the adjacency matrix. All within-frequency or amplitude-amplitude coupling networks are characterized by a symmetric adjacency matrix. However, to the best of our knowledge, no study until so far investigated the presence of a similar pattern in the adjacency matrix for CFS coupling which is strongly affected by the interactions due to higher harmonics of non-sinusoidal shape of the signals. The CFS adjacency matrix is by definition asymmetric. Actually, harmonic-driven spurious interactions result in symmetric CFS matrix. In other words, if the alpha activity in region *i* is coupled to the beta activity in region *j*, the (i,j)-th element of the adjacency matrix is non-zero. If this coupling is due to the non-sinusoidal shape of the waveform of the alpha-signals at both of these two regions, then the beta activity in region *i* is also synchronized to the alpha activity in region *j*, which results in a non-zero value at the (j, i)-th element of the adjacency matrix. This decreases the extent to which the adjacency matrix is asymmetric. Therefore, we reasoned that Harmoni should decrease the extent to which the adjacency matrix of the CFS network is symmetric. This idea was indeed confirmed as shown in figure 14-A with the boxplots of an asymmetry-index (refer to Methods) of the CFS networks before and after Harmoni for all subjects, where the asymmetry-index of the individual CFS connectivity networks increases significantly after Harmoni (two-sided paired t-test, *t*(80) ≈ 17.75, *p* ≈ 1.8*e* − 29). Furthermore, panel B of this figure shows that the percentage change of the asymmetry-index significantly decreases with the initial value of the index, pearson r= 0.75, *p* ≈ 0.0007 (with null hypothesis r=-0.55). In other words, Harmoni corrects the CFS network more (resulting in a more asymmetric network), when there are more potentially spurious interactions due to harmonics (i.e., the CFS network is less symmetric). See Methods for the rigorous statistical treatment of this analysis. Note that not all the harmonic-driven cross-frequency interactions are reflected in the symmetry of the CFS network adjacency matrix.

**Figure 14:**
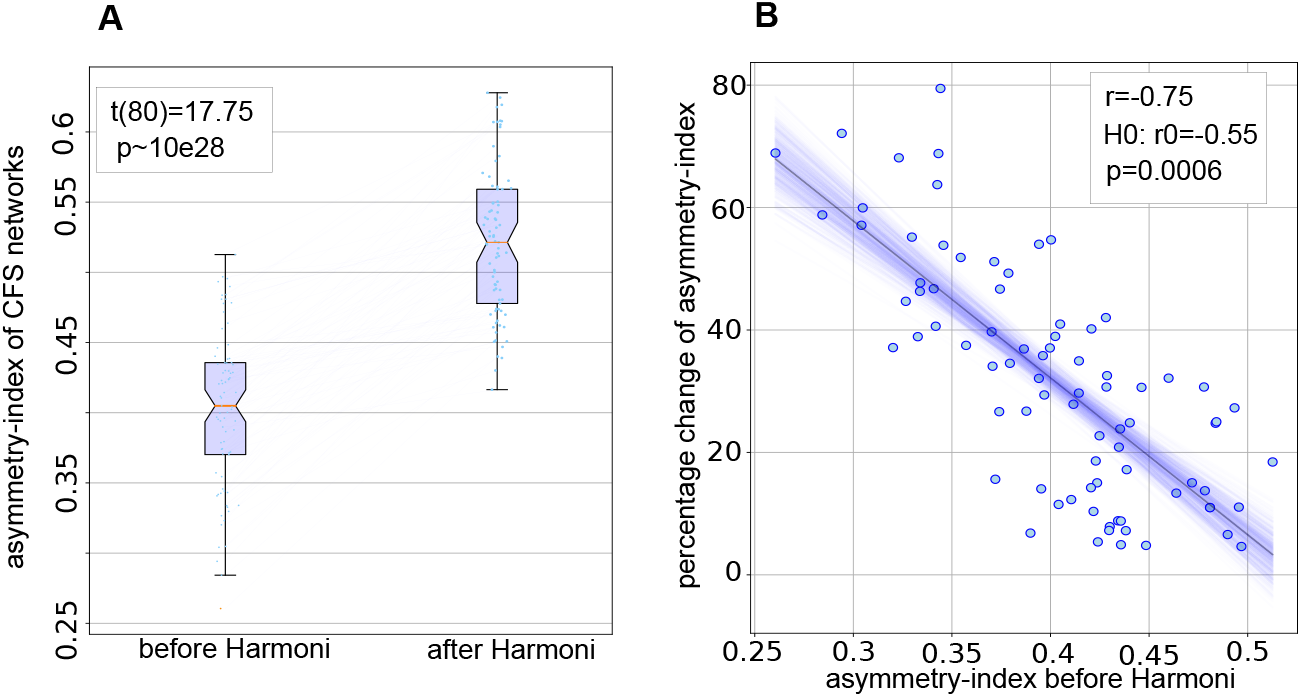
The CFS networks of individual LEMON subjects becomes more asymmetric after Harmoni. (A) the boxplots of the asymmetry-index of the CFS adjacency matrices of all subjects shows that the asymmetry of the CFS adjacency matrices increases significantly after Harmoni. (B) The scatter-plot of the percentage change of the asymmetry-index vs. the initial value of the index, i.e., before Harmoni.The less asymmetric the CFS network (i.e., the more harmonic-driven symmetric connections), the more changes are observed after Harmoni. The solid line shows the linear regression line and the blue shade shows the result of a leave-one-out bootstrap.

**Figure 15:**
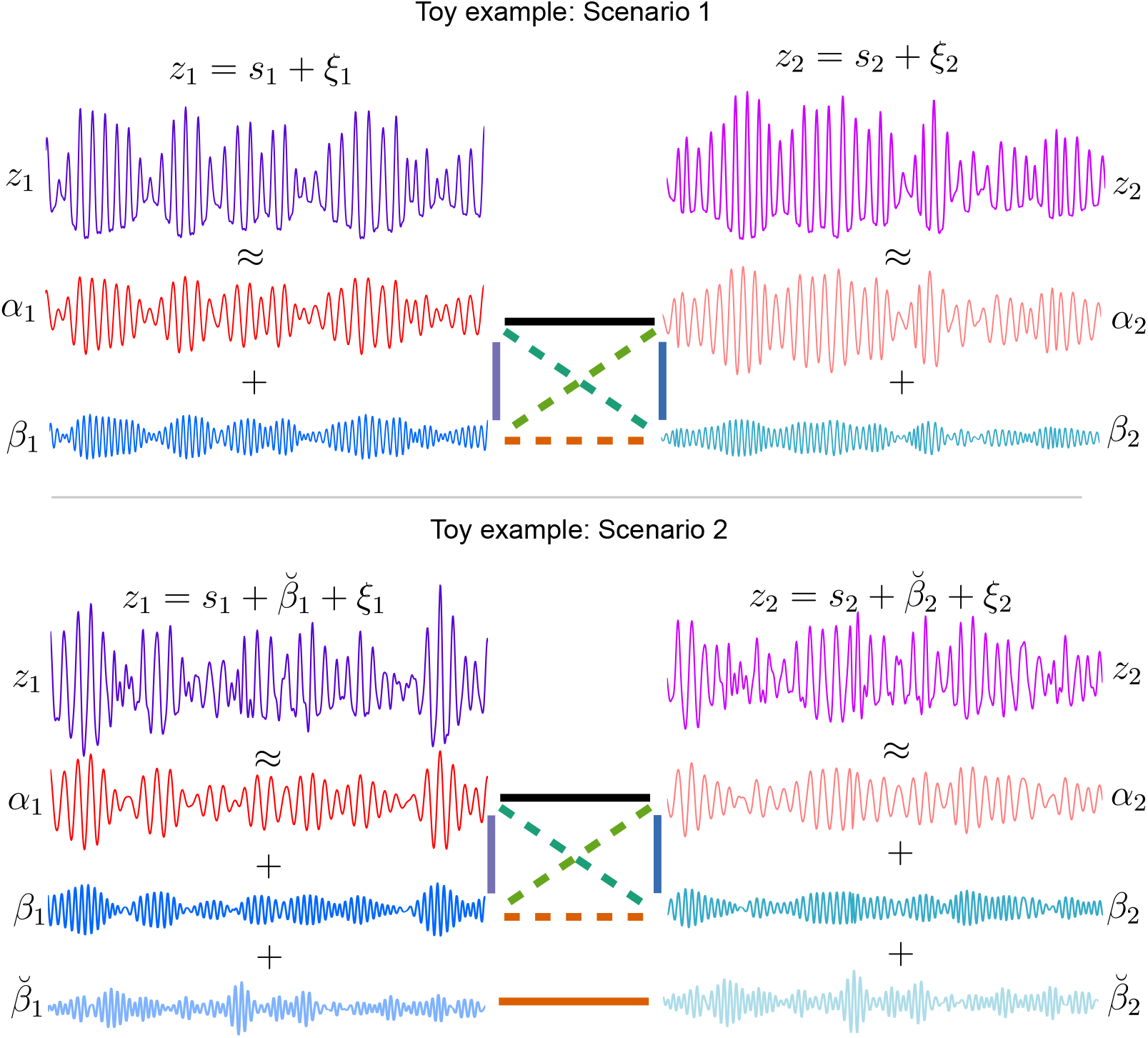
Examples of the composition of the two signals of scenario 1 and 2 of the toy examples. In scenario 1 *z_k_* = *s_k_* + *ξ_k_, k* = 1, 2 with *s_k_* = *α_k_* + *β_k_*. the noise components *ξ_k_* are not depicted. The solid lines show the simulated interactions, while the dashed lines are the spurious interactions which are the by-products of the simulated ones.

## 4. Discussion

EEG and MEG techniques are becoming more and more frequently used for the investigation of neuronal connectivity, owing to (1) their ability to record neuronal activity directly, and 2) their refined temporal resolution in a millisecond range which is required for the detection of subtle changes in neuronal dynamics. In addition, the recent advancement of brain data analysis for mapping sensor recordings to the cortex has provided an opportunity for computing the connectivity of different brain areas in source space. Yet, connectivity analysis with MEG/EEG faces considerable challenges. The limited spatial resolution and spatial mixing of neural activity from different regions hampers connectivity analysis. Additionally, the non-sinusoidal shape of brain oscillations has been repeatedly highlighted as crucially affecting the (mis)interpretation of underlying neuronal activity (Hyafil, 2017; Lozano-Soldevilla, 2018). Because non-sinusoidality always implies a presence of harmonics, these harmonics can often be mistakenly taken to represent genuine neuronal oscillations. Consequently, spurious interactions are observed between harmonics of a non-sinusoidal oscillation and other neuronal processes in the same frequency range, which in turn cannot be easily disentangled from genuine interactions. This has been recognized earlier as a major challenge for studying phase-amplitude coupling (PAC) in neuronal data (Aru et al., 2015; Giehl et al., 2021; Jensen et al., 2016; Lozano-Soldevilla et al., 2016; Zhang et al., 2021) as well as for n:m phase-synchronization (Hyafil, 2017; Scheffer-Teixeira and Tort, 2016; Siebenhühner et al., 2020). In this work, we directly addressed the issue of spurious interactions due to waveshape of oscillations and offer a solution for the assessment of phase synchronization as one of the most important measures used for connectivity analyses with brain electrophysiology (Marzetti et al., 2019; Nentwich et al., 2020; Sadaghiani et al., 2021; Vidaurre et al., 2020).

Currently available measures for quantifying n:m phase-synchronization (also referred to as cross-frequency synchronization - CFS) are not suitable for differentiation between genuine and spurious interactions. Short data length, filtering bias, and non-sinusoidal signal waveshape are being mentioned as reasons for measuring spurious n:m phase-synchronization. Statistical tests based on surrogate data can be used for disentangling spurious and genuine phase-synchronization due to limited data points or filtering factor. Yet, these procedures cannot differentiate the genuine interactions from the spurious ones due to the non-sinusoidality of oscillations (Scheffer-Teixeira and Tort, 2016). The reason for this is that Fourier and narrow-band analysis is the base of almost all current signal processing pipelines, where a signal is decomposed into narrow frequency band components. Consequently, the higher harmonics of a non-sinusoidal signal are analysed as representing genuine oscillations not directly relating to the fundamental frequency. In the context of cross-frequency coupling, this can result in the observation of spurious interactions which are mimicking genuine interactions and cannot be detected by surrogate tests. Furthermore, the non-sinusoidal waveshape of oscillatory brain signals produce spurious interactions in the within-frequency phase-synchronization in the range of harmonic-frequency, as depicted schematically in figure 1.

Although the presence of spurious interactions in phase-synchronization connectivity analysis of neurophysiological data has been largely acknowledged by the community, there has been only very few attempts for providing a potential solution for it. Palva et al. (2005) used the coincidence of cross-frequency phase-phase and amplitude-amplitude coupling as the hallmark of harmonic-driven CFS. This, however, is more a qualitative measure rather than a quantitative one and can be less applicable to the inter-areal whole brain connectivity analysis. In a recent paper, Siebenhühner et al. (2020) suggested a graph-theoretical analysis for discarding potential spurious CFS. The authors employed a procedure of detecting ambiguous motifs in the CFS graph combined with the within-frequency graphs of the fundamental and harmonic frequencies of interest, and discarding the CFS interactions corresponding to the links included in those motifs. This procedure, however, was not validated using realistic MEG/EEG simulations. Such graph-based post-processing of connectivity networks can in fact discard all the interactions which mimic the motif of spurious interactions in the connectivity graphs. However, due to the limited spatial resolution of MEG/EEG data, some of the genuine interactions among the ROIs may still coincide with harmonic-driven spurious interactions, as we show in figure 8-D. The graph motif of such interactions is similar to the spurious interactions, depicted in figure 8-A. Thus, a motif-discarding approach cannot distinguish the two cases of 8-A and D and would label the CFS interaction as a spurious one. Moreover, this graph-based correction method is applicable only to cross-frequency graphs, while, as discussed in this study, the within-frequency interactions in the harmonic frequency band may also include spurious interactions driven by non-sinusoidal waveshape. Therefore, to the best of our knowledge, so far there has been no method that can address the issue of spurious n:m interactions due to waveshape via removing the harmonic components from the neuronal signals.

### A signal processing tool for dealing with harmonics in connectivity

In this manuscript, we introduced the first signal processing tool for suppressing spurious within- and cross-frequency synchronization due to non-sinusoidal shape of the oscillatory activity in the brain. Our method significantly suppresses the spurious interactions, while at the same time not affecting genuine interactions present in data. We first validated these two key properties using simple, yet informing, simulations. They consisted of two signals with different components interacting with each other, giving us a chance to evaluate Harmoni’s performance in the presence of genuine and spurious interactions in data. The results of these simulations (figure 8) showed that Harmoni effectively suppresses spurious within- and cross-frequency interactions. Importantly, this suppression did not affect the genuine interactions.

### Realistic simulations: decrease in FPR, increase in AUC of ROC curve

In order to comprehensively assess Harmoni’s performance, we used realistic simulations where source mixing and limitations of source reconstruction are present. Using the area under curve (AUC) of the receiver operating characteristic (ROC) curve (figure 10), we showed that Harmoni increases the AUC of ROC curve of connectivity networks where the ground truth included both genuine and spurious interactions. This means that with Harmoni, it was possible to uncover even weak connections that would have been masked by spurious CFS otherwise. In the same direction as the results of the toy examples, the increase in AUC of ROC curve in realistic simulations indicates that Harmoni does not affect genuine interactions (reflected in TPR) and suppresses spurious interactions (i.e., false positives). In those simulations where the ground truth connectivity networks were based on spurious interactions only, Harmoni decreased the AUC of the FPR curve. Confirming other results of the simulations, this result further demonstrates that spurious interactions both for within-frequency and cross-frequency connectivity are indeed suppressed significantly by Harmoni. This aspect of Harmoni is particularly important for the investigation of connectivity for beta oscillations in the sensorimotor networks where comb-shaped mu oscillations are abundant (Schaworonkow and Nikulin, 2019) and thus their harmonics in beta frequency range should lead to spurious connectivity while merely reflecting interactions at the base alpha frequency. Additionally, in studies addressing the relationship of EEG and fMRI data, for example (Ritter et al., 2009), Harmoni could contribute to the suppression of the effects of harmonic components and disentangling the effect of harmonics and the genuine activity in the same frequency band.

Moreover, given that our simulations were based on hundreds of runs with different random locations of the sources, one can conclude that Harmoni is applicable to a wide variety of source configurations in the cortex including frontal, sensorimotor, and occipito-parietal areas.

### Harmoni on resting-state EEG data

Real neuronal data are of a complex nature and in most cases the ground truth of connectivity patterns is not known. Therefore, the main validating stage of new methods is rather based on simulations. However, any new method should also be applied to real data to further extend its validity. For this purpose, we used resting-state EEG (rsEEG) of 81 subjects from the LEMON database (Babayan et al., 2019). We discussed how a symmetric adjacency matrix of a cross-frequency synchronization network can reflect the presence of harmonics, and showed that the adjacency matrices of the CFS networks become more asymmetric after Harmoni. Additionally, we showed that Harmoni does not create new connections which were not observed before the application of Harmoni. However, it changes the relative strength of the already existing connections by suppressing spurious connectivity. Harmoni suppresses the CFS interactions both within and between regions, as depicted in figure 12-E. Consequently, other interactions, which were previously not ranked high due to the presence of strong spurious interactions, become more pronounced after the application of Harmoni. Although a detailed analysis of connectivity patterns of rsEEG goes beyond the scope of the current study, below we illustrate a few examples of the unmasked synchronization after the application of Harmoni.

In our data, only after the application of Harmoni, the visual cortical areas appear to be interacting strongly with other regions, especially inter-hemispherically. This in turn indicates that the interaction of the visual system with other cortical areas is not based only on a relatively slow amplitude-amplitude coupling as shown previously (Hipp and Siegel, 2015) but in fact can demonstrate genuine millisecond-range functional interactions important for the precise coordination of neuronal activity in the brain. Additionally, Wang et al. (2008), in an resting-state fMRI study, found that the spontaneous activity in primary visual cortex is associated with the activity in bilateral middle occipital gyrus, bilateral lingual gyrus, and bilateral cuneous and precuneus suggesting that these spontaneous activities may be related to visual imagery during resting-state. In our rsEEG data, the recovered inter-hemispheric interactions between the visual networks after the application of Harmoni can also be interpreted in this direction. Interestingly, figure 12-D shows the influence of Harmoni in recovering remote interactions of alpha and beta activity in ROIs overlapping with precuneus in both hemispheres - precuneus is known as a critical region for visual imagery in memory recall (Wang et al., 2008). Note that we also observed the emergence of precuneus as an important region in cross-frequency interactions, as well as in the inter-hemispheric interactions of visual cortices in our previous study (Idaji et al., 2020) with similar data, where phase-phase synchronized sources were separated with a multivariate source separation method.

Furthermore, figure 12-D illustrates intensified within- and inter-hemispheric interactions of default mode network (DMN) and visual networks, especially areas in the vicinity of V1. In line with our observation, in a recent paper, Costumero et al. (2020) reported a connectivity of V1 with DMN as well as posterior cingulate cortex in closed-eyes resting-state fMRI functional connectivity, suggesting that this connectivity may reflect a brain configuration associated with mental imagery.

### Harmoni and signal mixing

Due to the limited spatial resolution of non-invasive recordings, the activity of very close neuronal sources cannot be disentangled when being recorded by non-invasive imaging techniques such as MEG/EEG. Therefore, even at the source space, the observation of signals with non-sinusoidal shapes in non-invasive recordings may be due to mixing of distinct coupled sources with very close spatial locations. Using MEG/EEG, such cases cannot be distinguished from single sources generating signals with non-sinusoidal shapes. This limitation is also applicable to the Harmoni connectivity pipeline, when applying it to MEG/EEG data. However, it is important to note that, this problem is not a natural limitation of Harmoni. If we have access to invasive LFP recordings where the spatial resolution can be in the order of hundred of micrometers (Buzsáki et al., 2012), Harmoni can successfully resolve such cases.

The other aspect of spatial mixing relates to the leakage of spatially distanced source signals to other locations, even after source reconstruction. As a result, the synchronization observed at a single region (or even at a given reconstructed cortical source) may be due to the synchronization between distanced source signals which are spatially mixed and still could not be fully disentangled with source separation or source reconstruction methods. This, however, is again a general problem of data analysis in MEG/EEG research and is not specific to Harmoni. Therefore, in some instances the removal of harmonics in a ROI by Harmoni can lead to removing components which were not a harmonic of a lower frequency in that region but rather represents a leaked oscillatory activity from another coupled source. Yet, this property can in fact be an advantage for Harmoni: It can remove some of the spurious interactions which were present due to spatial leakage and uncover the activity at the harmonic frequency, which was not a result of spatial leakage of a coupled source. As an illustrative example for this property, in panel A of figure 8, if *β*_1_ is not a harmonic of *α*_1_ but a leakage of a crossfrequency coupled source different from *s*_1_, then the observed interaction *β*_1_ − *α*_2_ would still be accounted as a spurious interaction. This interaction, however, is successfully suppressed by Harmoni.

Finally, the mixing of background neuronal activity - known as 1/f noise - and other noise sources with oscillatory activities affects the signal-to-noise ratio (SNR) and consequently the estimation of the true phase of the oscillations. Using simulations in (Idaji et al., 2020), we showed how source separation of cross-frequency coupled sources worsens with decreasing SNR. Therefore, the phase estimates and consequently the n:m synchronization suffer from noise contamination. Because of this issue, the synchronization should be estimated with a sufficient amount of data points for MEG/EEG recordings.

## 5. Code and data availability

The codes of Harmoni, simulating toy examples, as well as analysing the simulated EEG and real data are available at github.com/harmonic-minimization. EEG data is from LEMON dataset, which is a public database (Babayan et al., 2019).

## 6. Author contributions

MJI: Conceptualization, Methodology, Software, Validation, Investigation, Formal analysis, Visualization, Project Administration, Writing - original draft, Writing - review editing. JZ and TS: Software, Writing - review editing. GN: Methodology, Writing - review editing. KRM: Writing - review editing, Supervision. AV: Resources, Writing - review editing, Supervision. VVN: Conceptualization, Methodology, Investigation, Project Administration, Writing - review editing, Supervision.

## 7. Competing interests

We declare no competing interests.

---

**Algorithm 1:**
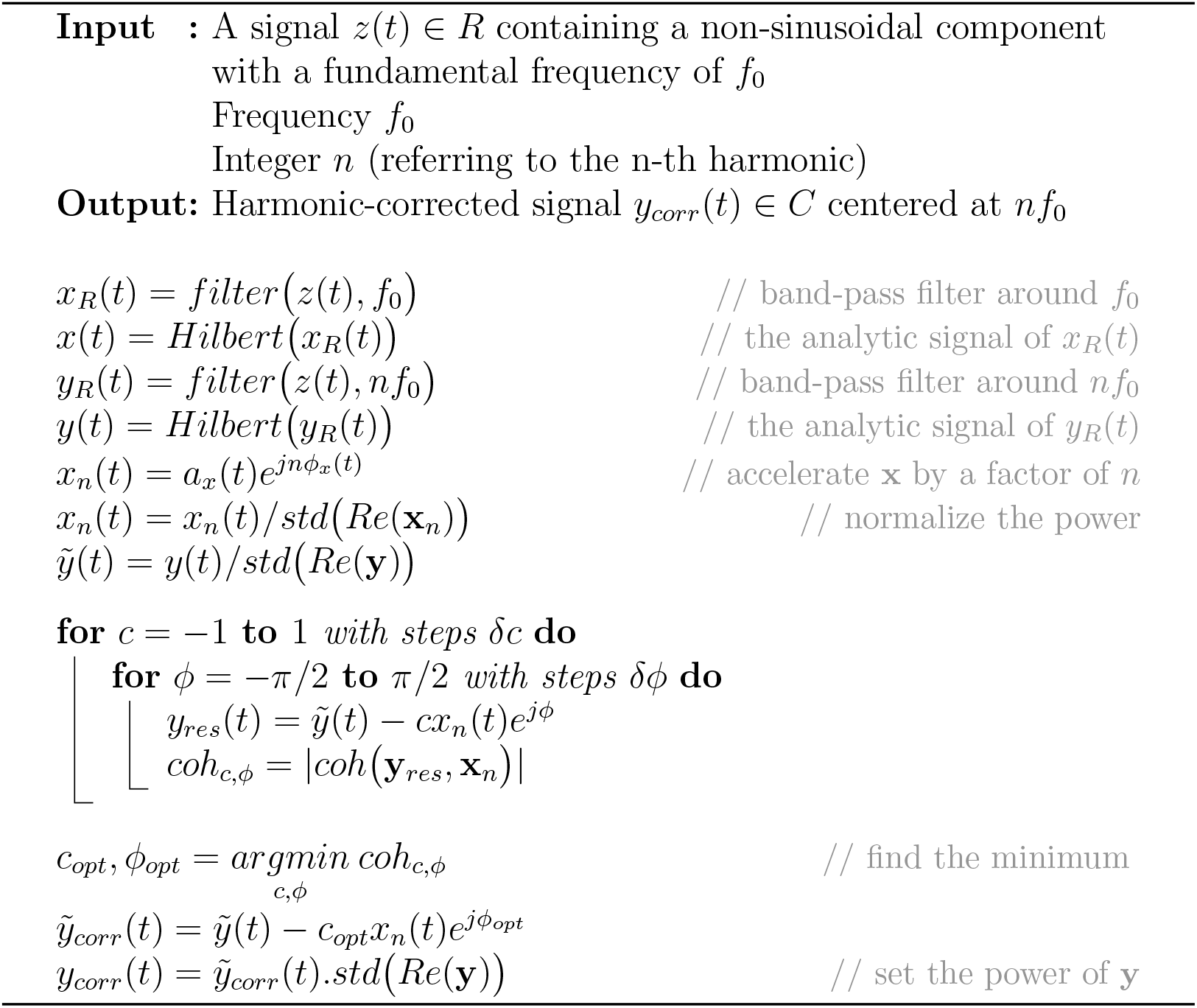
Grid-search algorithm of Harmoni. *filter*(., *f*_0_) stands for band-pass filtering around *f*_0_. *Hilbert*(.) builds the analytic signal of its input using the Hilbert transform. *Re*(.) denotes the real part of a complex number. *std*(.) stands for standard deviation.

## Notes

https://ftp.gwdg.de/pub/misc/MPI-Leipzig_Mind-Brain-Body-LEMON/EEG_MPILMBB_LEMON/EEG_Raw_BIDS_ID/

### Competing Interest Statement

The authors have declared no competing interest.

## References

Aru, J., Aru, J., Priesemann, V., Wibral, M., Lana, L., Pipa, G., Singer, W., Vicente, R., 2015. Untangling cross-frequency coupling in neuroscience. Current Opinion in Neurobiology 31, 51–61. doi:10.1016/j.conb.2014.08.002.

Babayan, A., Erbey, M., Kumral, D., Reinelt, J.D., Reiter, A.M., Röbbig, J., Schaare, H.L., Uhlig, M., Anwander, A., Bazin, P.L., et al., 2019. A mind-brain-body dataset of mri, eeg, cognition, emotion, and peripheral physiology in young and old adults. Scientific data 6, 1–21. doi:10.1038/sdata.2018.308.

Bullmore, E., Sporns, O., 2009. Complex brain networks: graph theoretical analysis of structural and functional systems. Nature reviews neuroscience 10, 186–198. doi:10.1038/nrn2575.

Buzsáki, G., Anastassiou, C.A., Koch, C., 2012. The origin of extracellular fields and currents—eeg, ecog, lfp and spikes. Nature reviews neuroscience 13, 407–420. doi:10.1038/nrn2575.

Buzsáki, G., Draguhn, A., 2004. Neuronal oscillations in cortical networks. science 304, 1926–1929. doi:10.1126/science.1099745.

Canolty, R.T., Knight, R.T., 2010. The functional role of cross-frequency coupling. Trends in cognitive sciences 14, 506–515. doi:10.1016/j.tics.2010.09.001.

Cole, S.R., Voytek, B., 2017. Brain oscillations and the importance of waveform shape. Trends in cognitive sciences 21, 137–149. doi:10.1016/j.tics.2016.12.008.

Costumero, V., Bueichekú, E., Adrian-Ventura, J., Ávila, C., 2020. Opening or closing eyes at rest modulates the functional connectivity of v1 with default and salience networks. Scientific reports 10, 1–10. doi:10.1038/s41598-020-66100-y.

Desikan, R.S., Ségonne, F., Fischl, B., Quinn, B.T., Dickerson, B.C., Blacker, D., Buckner, R.L., Dale, A.M., Maguire, R.P., Hyman, B.T., et al., 2006. An automated labeling system for subdividing the human cerebral cortex on mri scans into gyral based regions of interest. Neuroimage 31, 968–980. doi:10.1016/j.neuroimage.2006.01.021.

Engel, A.K., Fries, P., 2010. Beta-band oscillations—signalling the status quo? Current opinion in neurobiology 20, 156–165. doi:10.1016/j.conb.2010.02.015.

Fries, P., 2015. Rhythms for cognition: communication through coherence. Neuron 88, 220–235. doi:10.1016/j.neuron.2015.09.034.

Giehl, J., Noury, N., Siegel, M., 2021. Dissociating harmonic and non-harmonic phase-amplitude coupling in the human brain. NeuroImage 227, 117648. doi:10.1016/j.neuroimage.2020.117648.

Gramfort, A., Luessi, M., Larson, E., Engemann, D., Strohmeier, D., Brodbeck, C., Goj, R., Jas, M., Brooks, T., Parkkonen, L., Hämalainen, M., 2013. Meg and eeg data analysis with mne-python. Frontiers in Neuroscience 7, 267. doi:10.3389/fnins.2013.00267.

Gramfort, A., Luessi, M., Larson, E., Engemann, D.A., Strohmeier, D., Brodbeck, C., Parkkonen, L., Hämaläinen, M.S., 2014. Mne software for processing meg and eeg data. NeuroImage 86, 446 – 460. doi:10.1016/j.neuroimage.2013.10.027.

Harris, A.Z., Gordon, J.A., 2015. Long-range neural synchrony in behavior. Annual review of neuroscience 38, 171–194. doi:10.1146/annurev-neuro-071714-034111.

Hipp, J., Siegel, M., 2015. Bold fmri correlation reflects frequency-specific neuronal correlation. Current Biology 25, 1368–1374. doi:10.1016/j.cub.2015.03.049.

Hyafil, A., 2017. Disharmony in neural oscillations. Journal of Neurophysiology 118, 1–3. doi:10.1152/jn.00026.2017.

Idaji, M.J., Müller, K.R., Nolte, G., Maess, B., Villringer, A., Nikulin, V.V., 2020. Nonlinear interaction decomposition (nid): A method for separation of cross-frequency coupled sources in human brain. NeuroImage 211, 116599. doi:10.1016/j.neuroimage.2020.116599.

Jensen, O., Colgin, L.L., 2007. Cross-frequency coupling between neuronal oscillations. Trends in Cognitive Sciences 11, 267–269. doi:10.1016/j.tics.2007.05.003.

Jensen, O., Spaak, E., Park, H., 2016. Discriminating valid from spurious indices of phase-amplitude coupling. Eneuro 3. doi:10.1523/ENEURO.0334-16.2016.

Jones, S.R., Pritchett, D.L., Sikora, M.A., Stufflebeam, S.M., Hämälainen, M., Moore, C.I., 2009. Quantitative analysis and biophysically realistic neural modeling of the meg mu rhythm: rhythmogenesis and modulation of sensory-evoked responses. Journal of neurophysiology 102, 3554–3572. doi:10.1152/jn.00535.2009.

Kramer, M.A., Tort, A.B., Kopell, N.J., 2008. Sharp edge artifacts and spurious coupling in eeg frequency comodulation measures. Journal of neuroscience methods 170, 352–357. doi:10.1016/j.jneumeth.2008.01.020.

Lee, T.W., Girolami, M., Sejnowski, T.J., 1999. Independent component analysis using an extended infomax algorithm for mixed subgaussian and supergaussian sources. Neural computation 11, 417–441.

Lozano-Soldevilla, D., 2018. Nonsinusoidal neuronal oscillations: bug or feature? Journal of neurophysiology 119, 1595–1598. doi:10.1152/jn.00744.2017.

Lozano-Soldevilla, D., Ter Huurne, N., Oostenveld, R., 2016. Neuronal oscillations with non-sinusoidal morphology produce spurious phase-to-amplitude coupling and directionality. Frontiers in computational neuroscience 10, 87. doi:10.3389/fncom.2016.00087.

Marzetti, L., Basti, A., Chella, F., D’Andrea, A., Syrjäla, J., Pizzella, V., 2019. Brain functional connectivity through phase coupling of neuronal oscillations: A perspective from magnetoencephalography. Frontiers in Neuroscience 13, 964. doi:10.3389/fnins.2019.00964.

Miller, K.J., Schalk, G., Fetz, E.E., den Nijs, M., Ojemann, J.G., Rao, R.P., 2010. Cortical activity during motor execution, motor imagery, and imagery-based online feedback. Proceedings of the National Academy of Sciences 107, 4430–4435. doi:10.1073/pnas.0913697107.

Nentwich, M., Ai, L., Madsen, J., Telesford, Q.K., Haufe, S., Milham, M.P., Parra, L.C., 2020. Functional connectivity of eeg is subject-specific, associated with phenotype, and different from fmri. NeuroImage 218, 117001. doi:10.1016/j.neuroimage.2020.117001.

Nikulin, V.V., Brismar, T., 2006. Phase synchronization between alpha and beta oscillations in the human electroencephalogram. Neuroscience 137, 647–657. doi:10.1016/j.neuroscience.2005.10.031.

Nolte, G., Bai, O., Wheaton, L., Mari, Z., Vorbach, S., Hallett, M., 2004. Identifying true brain interaction from eeg data using the imaginary part of coherency. Clinical neurophysiology 115, 2292–2307. doi:10.1016/j.clinph.2004.04.029.

Palva, J.M., Palva, S., 2018a. Functional integration across oscillation frequencies by cross-frequency phase synchronization. European Journal of Neuroscience 48, 2399–2406. doi:10.1111/ejn.13767.

Palva, J.M., Palva, S., Kaila, K., 2005. Phase synchrony among neuronal oscillations in the human cortex. Journal of Neuroscience 25, 3962–3972. doi:10.1523/JNEUROSCI.4250-04.2005.

Palva, S., Palva, J.M., 2018b. Roles of brain criticality and multiscale oscillations in temporal predictions for sensorimotor processing. Trends in neurosciences 41, 729–743. doi:10.1016/j.tins.2018.08.008.

Pascual-Marqui, R.D., 2007. Discrete, 3D distributed, linear imaging methods of electric neuronal activity. Part 1: exact, zero error localization. arXiv preprint arXiv:0710.3341.

Patzelt, F., 2019. colornoise python package. https://pypi.org/project/colorednoise/. Checked: 2020-06-24.

Ritter, P., Moosmann, M., Villringer, A., 2009. Rolandic alpha and beta eeg rhythms’ strengths are inversely related to fmri-bold signal in primary somatosensory and motor cortex. Human brain mapping 30, 1168–1187. doi:10.1002/hbm.20585.

Sadaghiani, S., Brookes, M.J., Baillet, S., 2021. Connectomics of human electrophysiology. PsyArXiv.

Sadaghiani, S., Kleinschmidt, A., 2016. Brain networks and *α*-oscillations: structural and functional foundations of cognitive control. Trends in cognitive sciences 20, 805–817. doi:10.1016/j.tics.2016.09.004.

Schaefer, A., Kong, R., Gordon, E.M., Laumann, T.O., Zuo, X.N., Holmes, A.J., Eickhoff, S.B., Yeo, B.T., 2018. Local-global parcellation of the human cerebral cortex from intrinsic functional connectivity mri. Cerebral cortex 28, 3095–3114. doi:10.1093/cercor/bhx179.

Schaworonkow, N., Nikulin, V.V., 2019. Spatial neuronal synchronization and the waveform of oscillations: Implications for eeg and meg. PLoS Computational Biology 15, e1007055. doi:10.1371/journal.pcbi.1007055.

Scheffer-Teixeira, R., Tort, A.B., 2016. On cross-frequency phase-phase coupling between theta and gamma oscillations in the hippocampus. Elife 5, e20515. doi:10.7554/eLife.20515.

Siebenhühner, F., Wang, S.H., Arnulfo, G., Lampinen, A., Nobili, L., Palva, J.M., Palva, S., 2020. Genuine cross-frequency coupling networks in human resting-state electrophysiological recordings. Plos Biology 18, e3000685. doi:10.1371/journal.pbio.3000685.

Tu, Y.K., 2016. Testing the relation between percentage change and baseline value. Scientific reports 6, 1–8. doi:10.1038/srep23247.

Vidaurre, C., Haufe, S., Jorajuría, T., Müller, K.R., Nikulin, V.V., 2020. Sensorimotor functional connectivity: a neurophysiological factor related to bci performance. Frontiers in Neuroscience 14, 1278. doi:10.3389/fnins.2020.575081.

Wang, K., Jiang, T., Yu, C., Tian, L., Li, J., Liu, Y., Zhou, Y., Xu, L., Song, M., Li, K., 2008. Spontaneous activity associated with primary visual cortex: a resting-state fmri study. Cerebral cortex 18, 697–704. doi:10.1093/cercor/bhm105.

Xia, M., Wang, J., He, Y., 2013. Brainnet viewer: a network visualization tool for human brain connectomics. PloS one 8, e68910. doi:10.1371/journal.pone.0068910.

Yeo, B.T., Krienen, F.M., Sepulcre, J., Sabuncu, M.R., Lashkari, D., Hollinshead, M., Roffman, J.L., Smoller, J.W., Zöllei, L., Polimeni, J.R., et al., 2011. The organization of the human cerebral cortex estimated by intrinsic functional connectivity. Journal of neurophysiology doi:10.1152/jn.00338.2011.

Zhang, J., Idaji, M.J., Villringer, A., Nikulin, V.V., 2021. Neuronal biomarkers of parkinson’s disease are present in healthy aging. NeuroImage 243, 118512. URL: https://www.sciencedirect.com/science/article/pii/S1053811921007850, doi:10.1016/j.neuroimage.2021.118512.

